# Dopamine neurons drive spatiotemporally heterogeneous striatal dopamine signals during learning

**DOI:** 10.1101/2023.07.01.547331

**Authors:** Liv Engel, Amy R Wolff, Madelyn Blake, Val L. Collins, Sonal Sinha, Benjamin T Saunders

**Author notes:** Correspondence: BTS (, @bensaunders, saunderslab.com). Denotes equal contribution (Liv Engel, Amy R Wolff).

## Abstract

Environmental cues, through Pavlovian learning, become conditioned stimuli that invigorate and guide animals toward acquisition of rewards. Dopamine neurons in the ventral tegmental area (VTA) and substantia nigra (SNC) are crucial for this process. Dopamine neurons are embedded in a reciprocally connected network with their striatal targets, the functional organization of which remains poorly understood. Here, we investigated how learning during optogenetic Pavlovian cue conditioning of VTA or SNC dopamine neurons directs cue-evoked behavior and shapes subregion-specific striatal dopamine dynamics. We used a fluorescent dopamine biosensor to monitor dopamine in the nucleus accumbens (NAc) core and shell, dorsomedial striatum (DMS), and dorsolateral striatum (DLS). We demonstrate spatially heterogeneous, learning-dependent dopamine changes across striatal regions. While VTA stimulation evoked robust dopamine release in NAc core, shell, and DMS, cues predictive of this activation preferentially recruited dopamine release in NAc core, starting early in training, and DMS, late in training. Corresponding negative prediction error signals, reflecting a violation in the expectation of dopamine neuron activation, only emerged in the NAc core and DMS, and not the shell. Despite development of vigorous movement late in training, conditioned dopamine signals did not similarly emerge in the DLS, even during Pavlovian conditioning with SNC dopamine neuron activation, which elicited robust DLS dopamine release. Together, our studies show broad dissociation in the fundamental prediction and reward-related information generated by different dopamine neuron populations and signaled by dopamine across the striatum. Further, they offer new insight into how larger-scale plasticity across the striatal network emerges during Pavlovian learning to coordinate behavior.

## INTRODUCTION

Behavior is heavily influenced by external sensory information (“cues”) that acquires motivational value to trigger and guide complex behaviors. Midbrain dopamine neuron activity, and dopamine signals in the striatum, are crucial for learning and evaluation of cues, actions, and biologically relevant outcomes during decision making (1–5). Dopamine neuron activity is evoked by rewards and can substitute for rewards to drive cue-based learning and motivation, with heterogeneous functions ascribed to different subcircuits (6–13).

Dopamine neurons project largely along an ipsilateral, ventromedial to dorsolateral gradient, with ventral tegmental area (VTA) neurons preferentially innervating the ventral and medial striatum, and substantia nigra. (SNC) neurons preferentially innervating the dorsal and lateral striatum (14–16). Through direct and indirect circuits back from the striatum to the midbrain, this network is thought to be functionally organized as an ascending “spiral” (16,17), where the locus of control of learning and behavioral execution transitions from dopamine inputs into the nucleus accumbens (NAc) to dopamine inputs into the dorsolateral striatum (DLS) (18–20). This transition is thought to parallel the emergence of stereotyped, habitual action patterns seen with extended training (21–23). Recent work has complicated this influential framework, however, with studies showing a lack of DLS recruitment (24,25), no reduction in NAc dopamine signaling (26,27), and a maintenance of NAc control of behavior late in conditioning (28). In parallel, there is growing consensus of highly specialized dopamine signaling patterns and functional roles among dopamine projections and the striatal niches they innervate (6,8,26,29–38).

Despite the implications for a more complete understanding of the functional architecture of the striatal network, it remains unknown how dopamine neuron activity itself engages dopamine signals broadly throughout the striatum, across different phases of learning. To establish the impact of dopamine neuron activity during Pavlovian learning on cue and movement related dopamine signaling, we combined optogenetic manipulation of VTA or SNC dopamine neurons with simultaneous recording of dopamine transmission in different striatal subregions, using the biosensor dLight (39), and pose estimation (40) for detailed behavioral analysis. Brief dopamine neuron activation drove learning about antecedent cues, and these cues came to evoke conditioned approach and rapid movement invigoration, similar to learning driven by external rewards (6,41,42). Critically, striatal dopamine dynamics during dopamine-driven Pavlovian learning were spatiotemporally heterogeneous and qualitatively distinct for VTA versus SNC activation. VTA cue-evoked dopamine signals emerged early in training in the NAc core, but not NAc shell, and then later emerged in the dorsomedial striatum (DMS). NAc core and DMS dopamine signals were also selectively sensitive to prediction errors related to VTA dopamine neuron activity. Despite extended training that promoted vigorous VTA cue-evoked movement, NAc core signals did not diminish and cue-evoked signals in the DLS did not emerge. In contrast, SNC dopamine neuron predictive cues promoted robust conditioned behavior, but did not engage similar cue-evoked dopamine signals in the DLS or NAc core.

Our results indicate that Pavlovian learning mediated by dopamine neuron activity leads to a preferential recruitment of cue-, reward, and error-related dopamine signals in only a subset of striatal targets. Further, our data suggest that extended Pavlovian learning does not produce a ventral-to-dorsal or dorsal-to-ventral shift in dopamine signaling. Instead, different populations of dopamine neurons engage qualitatively distinct recruitment of striatal dopamine signaling during learning, across different timescales and with unique subregion-based profiles.

## RESULTS

### VTA dopamine neurons drive cue learning and vigorous behavior

For manipulation of dopamine neurons, we expressed the red-shifted excitatory opsin Chrimson in TH-cre rats (6,43), and implanted optic fibers over the center of the VTA (**Fig 1A,B; Supplemental Fig 2**). To investigate the ability of sensory cues paired with activation of VTA dopamine neurons to drive behavioral invigoration, we made use of an optogenetic Pavlovian conditioning procedure (6). Rats first underwent brief habituation to a novel, neutral cue (light+tone, 7 sec), followed by 20 daily sessions where the cue was paired with laser delivery to the VTA (589 nm, 20Hz, 2mW pulses, 5 sec) 25 times per session (**Fig 1C**). Laser stimulation began 2 sec after cue onset. No other reward or external stimuli were administered.

**Figure 1.**
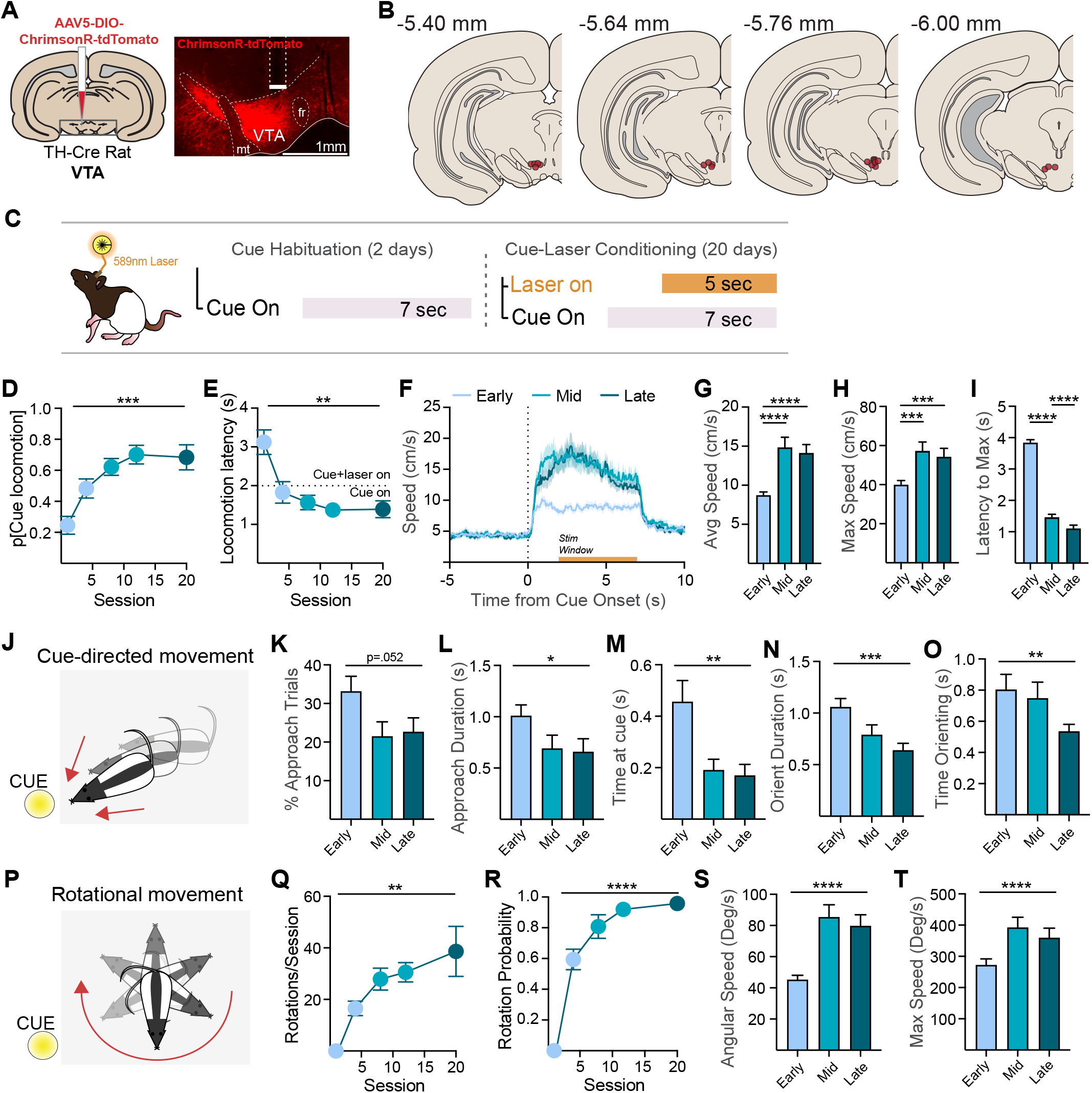
VTA dopamine neurons drive cue learning and vigorous behavior that evolves over time. A) ChrimsonR-tdTomato was expressed in dopamine neurons in TH-cre rats (N=20) and optic fibers were inserted over viral expression in the VTA. B) Location of fiber tips in the VTA. C) Schematic of optogenetic Pavlovian conditioning procedure. Following habituation to a novel, neutral cue (light + tone), rats underwent training where the cue was paired with laser (589 nm) delivery 25 times per session for 20 sessions. D) Across this training, conditioned responses (movement) emerged with high probability in the first 2-sec of the cue period. E) The latency of locomotion onset decreased across sessions and the majority of responses were initiated less than 2-sec after cue onset, indicating they were cue, rather than laser evoked. F) Movement was tightly locked to cue onset, increasing in speed across the early (days 1-2), middle (days 10-12), and late (days 19-20) phases of training. Both average G) and maximum H) speed achieved increased between early and middle/late training phases. I) The time required to reach maximum speed decreased across each training phase, indicating greater movement acceleration. J) We also assessed the path of movement patterns, finding that rats emitted conditioned behavioral responses directed at the cue. K) Further classification using semi-automated pose estimation data showed a trend towards a decrease in the percentage of trials containing an approach response across training. L) Approach duration and M) the time rats spent at the cue across all trials decreased across training phases. N) We also quantified cue orientation, measured by comparing the rat’s head direction to the cue position, finding that on trials where an orientation occurred, the length of orientations decreased and O) the amount of orientation across all trials decreased as training progressed. P) As training progressed, cue-evoked movement evolved into a more vigorous, rotational pattern, as rats ran in circles around the chamber. Rotation did not occur early in training but increased robustly in Q) number and R) probability across training. S) Correspondingly, the average and T) maximum angular speed during cue presentations increased robustly across training. ****p<.0001, ***p<.001, **p<.01, *p<.05.

We first analyzed a subset of video recordings of training sessions (n=10 rats) via experimenter inspection. Rats quickly exhibited conditioned behavioral responses (CRs), first defined simply as locomotion, which increased in probability across training during cue presentations (**Fig 1D**; probability of locomotion in first 2-sec of cue, mixed-effects ANOVA main effect of session, F(1.53,13.04)=15.75, p=.0006). Conditioned movement emerged at shorter onset latency as training progressed, with the large majority of CRs beginning during the first 2-sec of the cue, before laser onset (**Fig 1E**; F(1.84,11.93)=12.62, p=.0013). This is consistent with previous findings (6), indicating that CRs elicited by dopamine-paired cues reflect learned behavioral responses, and are not simply triggered by laser activation of dopamine neurons. We have previously demonstrated (6) that this conditioned invigoration is dependent on temporal contiguity between the cue and dopamine neuron activation: unpaired, uncued optogenetic activation of dopamine neurons in either the VTA or SNC is insufficient to drive cue learning, and does not evoke movement or other behaviors (27,47). For more detailed quantification of cue-evoked behavior across the whole dataset (n=20 rats), we employed DeepLab-Cut (40) for movement analysis (**Supplemental Fig 1**). This revealed that the latency of movement onset after cue onset decreased (mixed effects ANOVA main effect of training phase, F(1.7,28.8)=3.79, p=.041) and the duration of movement bouts during cue presentations increased (F(1.51,25.59)=22.04, p<.0001) across early (days 1-2), middle (days 10-12), and late (days 19-20) training phases. The average (**Fig 1F,G**) and maximum speed (**Fig 1H**) of conditioned movement both increased across training (F(1.81,30.81)=25.65, p<.0001; F(1.928,32.78)=21.52, p<.0001).

Further, the vigor of conditioned movement accelerated across training, reflected in a decreased latency to achieve maximum speed (**Fig 1I**; F(1.44,24.43)=327.9, reflected a conditioned, cue-dependent invigoration state, as examination of speed during the 5 seconds before cue onset show no changes across training (Average pre cue speed, no effect of training phase, F(1.99,33.9)=.036, p=.964); Max pre cue speed, no effect of training phase, F(1.92, 32.6)=1.44, p=.858).

We analyzed behavioral data split by both phase and sex (12 F, 8 M). Overall, female rats had a higher average cue-evoked movement speed than males at each training phase (**Supplementary Fig 3B**; 2-way mixed effects ANOVA main effect of sex, F(1,18)=4.42, p=.0499). This reflected a general sex difference in movement, rather than a sex-specific difference in dopamine-mediated learning, as females also had a higher pre-cue movement speed across all training phases (2-way mixed effects ANOVA main effect of sex, F(1,18)=4.65, p=.045). We found no other sex-based differences in behavioral measures (**Supplementary Fig 3**; all sex comparisons p>.05).

The visual component of the cue complex was a circular illuminated panel positioned at rearing height on one wall of the chamber. This allowed, for localized, cue-directed CRs to occur, in the form of orientation, approach, and investigation (**Fig 1J**), a common behavioral response to reward-predictive cues (48). We quantified cue approach behavior in two ways. First, experimenters recorded instances of approach. Approach behavior was evident early in training, on approximately one-third of cue presentations, and trended downward throughout conditioning (approach number: mixed effects ANOVA trend for training phase, F(2.79,23.73)=2.98, p = 0.0525; approach probability: F(2.65,22.55)=2.87, p = 0.064).

For characterization of detail of movement paths rats took in response to the cue, we used a semi-automated behavioral classification method, based on rat position and body posture data from the DeepLabCut model. Behavioral classification based on DLC position data was highly consistent with human-based video scoring but allowed for a more detailed assessment of cue-directed behavior. Automated detection of approach using the DLC-based positional location of the nose relative to the cue was classified as periods when the animals nose approached the cue for periods longer than 0.2s in duration. In line with our manually-scored data, this analysis showed that approach behavior developed early in Pavlovian training and trended downward throughout. There was a decreasing trend in the likelihood of approaches initiated after cue onset from early to late training days (**Fig 1K**, Percentage cue periods with an approach: mixed effects ANOVA main effect of training phase, F(1.716,29.17)=3.43, p = 0.052). Approach latency was stable across training (latency: F(1.67,26.78)=0.25, p=.74). On trials where there were approaches, however, rats spent less time engaged in cue approach from early to later phases of training (**Fig 1L**, approach duration: F(1.44,24.41)=11.07, p=.001) and the time spent interacting with the cue across all trials decreased from early to later training phases (**Fig 2M**, F(1.85,29.51)=5.00, p=.01).

**Figure 2.**
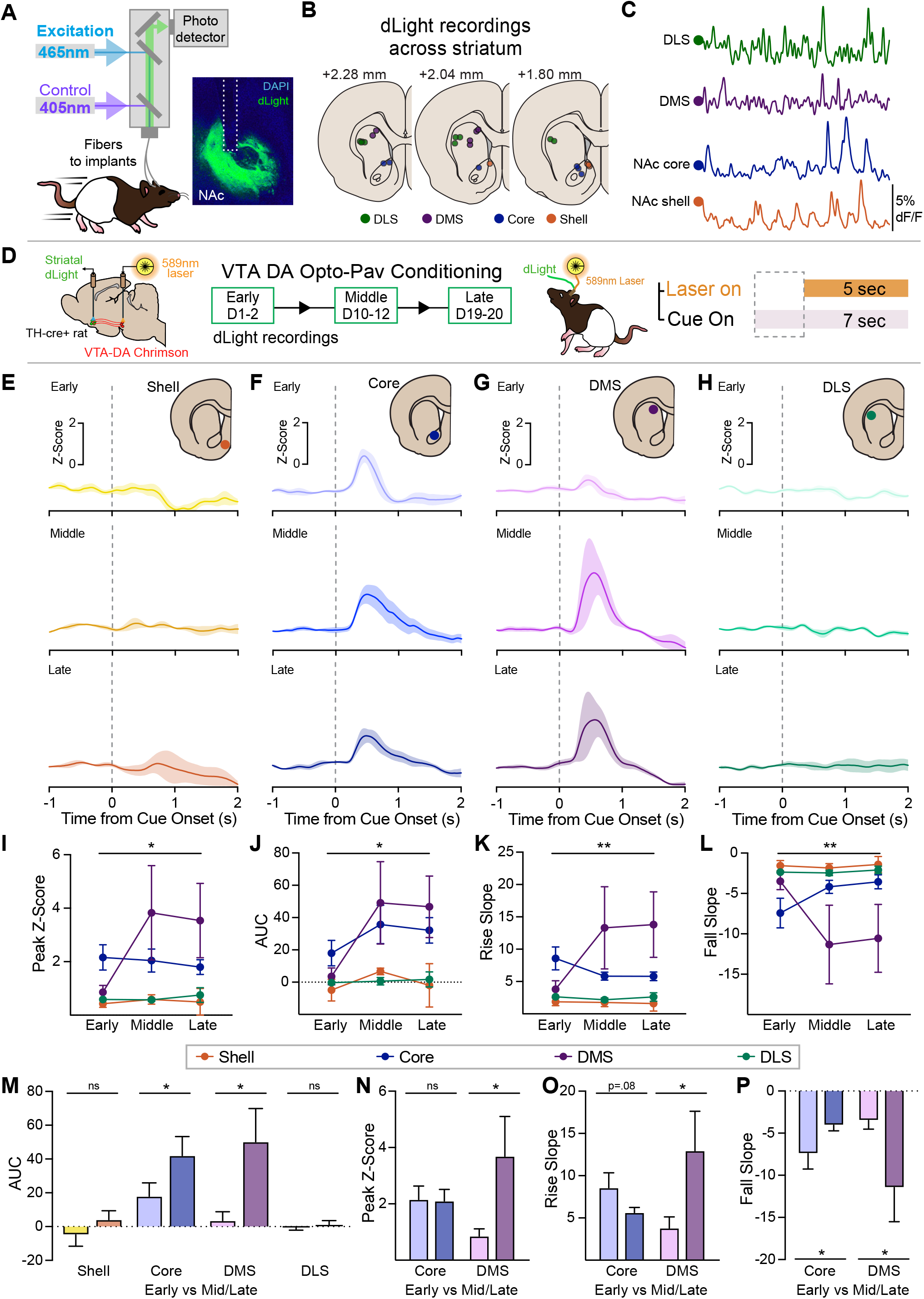
Spatially and temporally heterogeneous conditioned dopamine signals develop to VTA dopamine neuron predictive cues. A) Schematic of fiber photometry system and analysis. The dopamine biosensor dLight 1.3b was expressed in the striatum of TH-cre rats expressing ChrimsonR in VTA dopamine neurons. dLight signals were fitted against an isosbestic signal, to control for photobleaching and movement artifacts. B) Recording locations for photometry measurements made from four striatal regions: the NAc medial shell (n=4), NAc core (n=6), DMS (n=6), and DLS (n=9). C) Robust spontaneous dopamine fluctuations were identified in each region. D) Overview of recording scheme during behavioral testing. dLight signals were recorded across optogenetic Pavlovian conditioning in three training phases: early (days 1-2), middle (days 10-12), and late (days 19-20). Data in this figure correspond to the first 2-sec of the cue periods. E-H) Phasic dopamine signals in response to cues paired with optogenetic activation of VTA dopamine neurons developed only in some striatal targets. E) In the medial NAc shell, cue-evoked dopamine signals did not emerge at any training phase. F) In the NAc core, a phasic dopamine response emerged, which remained stable across training. G) In the DMS, no cue-evoked signals were present early in training, but they emerged at the middle and late phases. H) In the DLS, consistent cue-evoked signals did not emerge at any training phase. I-L) Cue onset-related signal dynamics, including signal peak, AUC, and the rise and fall slopes or peak signal, varied across region and training phase. M) AUC of cue-evoked signal increased only in the core and DMS regions from early to later in training. N) The peak Z-score increased in the DMS but not core. O) The rise slope of the cue-evoked signal increased in the DMS only. P) The fall slope of the peak signal decreased in the core and increased in the DMS. **p<.01, *p<.05.

We also assessed orientation to the cue, as a measure of cue-directed attention that did not meet the threshold of approach. We calculated head direction, based on interpolation of a vector projecting from the cranial implant to the nose, which allowed for quantification of head angle compared to other features of the environment (**Supplemental Fig 1**). Given that the cue light was in a fixed position relative to the rat, orientation behavior was interpreted as a reduction in head-to-cue angle (< 30°, 0.2s duration). The likelihood of orientation to the cue increased from early to later stages of training (Percentage of trials: mixed effects ANOVA main effect of training phase, F(1.476,25.08)=9.05 p<.0.01), with a consistent onset latency (F(1.71,29.14)=1.36, p=.27). However, on trials where animals did orient to the cue, the total time orienting (**Fig 1N**, F(1.98,33.64)=8.88, p=.0008) and the duration of orientation across all trials (**Fig 1O**, F(1.56,26.45)=5.53, p=.01) decreased significantly in the later stages of training. Thus, overall, rats maintained attention to the cue across training, but spent less time engaged in cue investigation.

As training progressed, we observed an evolution in conditioned behavior, where rats transitioned from pre-dominantly linear movement often directed at the cue, to more vigorous rotational behavior around the perimeter of the chamber (**Fig 1P, Supplementary Fig 1**). This turning behavior was cue-evoked, and always occurred contralateral to the stimulation site. Rotational behavior was quantified via video inspection. Rotations never occurred in the first few sessions of training, indicating they reflect a learned conditioned response rather than an automatic laser-evoked movement pattern. Across the middle and later training stages, rotational movement strongly emerged, increasing in number (**Fig 1Q**, Mixed effects ANOVA main effect of training phase, F(1.44,12.25)=12.83, p=.0019) and probability (**Fig 1R**, F(1.98,16.84)=77.39, p<.0001) during cue presentations. Using our DLC network, we next interpolated the rat’s angular speed as another indication of movement invigoration during cue presentations. We found that the average (**Fig 1S**, F(1.49,25.32)=30.38, p<.0001) and maximum (**Fig 1T**, F(1.52,25.83)=6.68, p=.008) angular speed increased significantly across training phases.

Together these results show that, via association with VTA dopamine neuron activation, cues acquired Pavlovian conditioned stimulus properties, driving robust conditioned behaviors that became more vigorous and less cue directed with extended dopamine-mediated learning.

### VTA dopamine mediated learning drives cue-evoked dopamine signals first in the core, then DMS, but not in shell or DLS

Cues paired with natural rewards evoke dopamine neuron activity and release at dopamine neuron targets (12,13,31,49). Here, we assessed the extent to which VTA dopamine-paired cues evoke dopamine signaling downstream in the striatum (**Fig 2**), using fiber photometry recordings of the dLight1.3b sensor (39). Dopamine signaling was measured across four striatal regions: the NAc medial shell, NAc core, DMS, and DLS (**Fig 2B**). Across all sites, we found robust spontaneous dopamine signals (**Fig 2C**). To further validate this approach, we include several analyses (**Supplemental Fig 4**) comparing the 465 and 405-nm signal channels across regions, showing robust signal-to-noise fidelity. Further, we find qualitatively similar patterns in the optogenetic conditioning data when the 465 channel signal was processed independent of 405-channel fitting, indicating that isosbestic fitting for delta F/F calculations in the subsequent analyses did not artificially create region-specific patterns of activity (50,51).

We recorded dLight signals at these sites across optogenetic Pavlovian conditioning of VTA dopamine neurons (**Fig 2D**). Spatially and temporally heterogeneous dopamine signals emerged in response to VTA dopamine neuron predictive cues (**Fig 2E-H**). Cue-evoked increases in dopamine developed in the NAc core and the DMS, but not in the medial shell or DLS. NAc core signals emerged first, early in training, and remained stable (**Fig 2F**). In contrast, DMS signals were not present early in training, but emerged by the middle phase (**Fig 2G**), once behavior had largely stabilized. For the core and DMS, dopamine signals peaked around 800 ms after cue onset, before laser stimulation began. Despite the development of rapid and vigorously expressed movement in response to the cue, consistent cue conditioned dopamine signals in the DLS did not emerge (**Fig 2H**). Qualitative differences in striatal dopamine dynamics were apparent from examination of average traces, but we also quantified several signal metrics during the first 2-sec of cue presentations that differed across regions and training phases, including the cue-evoked peak Z-score (**Fig 2I**, mixed effects ANOVA interaction of region and phase, F(6,38)=3.14, p=.013; main effect of region F(3,21)=4.43, p=.015), area under the curve (**Fig 2J**, interaction of region and phase, F(6,38)=2.56, p=.035; main effect of region, F(3,21)=5.26, p=.0073), and the rise (**Fig 2K**, interaction of region and phase, F(6,38)=4.11, p=.0033; main effect of region, F(3,21)=4.60, p=.013) and fall slopes (**Fig 2L**, interaction of region and phase, F(6,37)=4.00, p=.0035; main effect of region, F(3,21)=3.99, p=.022) of the peak signal.

Average NAc shell and DLS cue-related signal AUCs remained near zero and did not change across training (**Fig 2M**, paired t test for shell, t(3)=1.0, p=.39; paired t test for DLS, t(8)=.47, p=.65). In the core, while the peak Z-score (**Fig 2N**, t(5)=.09, p=.48) did not change, the AUC increased (**Fig 2M**; t(5)=2.05, p=.048) and the fall slope decreased (**Fig 2P**, t(5)=2.23, p=.038), suggesting more prolonged cue-evoked dopamine release, or a change in dopamine reuptake rate, across training. In the DMS, the AUC (**Fig 2M**, t(5)=2.55, p=.026) and peak (**Fig 2N**, t(5)=2.18, p=.04) Z-score increased between the early and middle/late training phases, suggesting dopamine signals in this region were recruited only after initial learning. The rise (**Fig 2O**, t(5)=2.34, p=.033) and fall (**Fig 2P**, t(5)=2.04, p=.048) slopes of the DMS signal also increased, suggesting an increase in the rate of dopamine release and clearance as training progressed. We compared the within-session evolution of cue-evoked signals for the core and DMS, by separately analyzing the first 5 versus last 5 trials in the early and middle training phases (**Supplemental Fig 5**). This showed that NAc core cue signals were apparent immediately in the first session. DMS cue signals, in contrast, were not yet present by the end of the early training phase, emerging later, in the middle phase.

We next examined the relationship between cue-evoked dopamine signals and anatomical placement of recordings sites in the striatum, independent of regional subgrouping. This revealed separate gradients spanning the medial-lateral axis of the ventral versus dorsal striatum (**Supplemental Fig 6**). These gradients were opposing in their pattern, and developed on different timescales. In the ventral striatum (NAc core and medial shell placements), cue-evoked dopamine signals were smallest in more medially positioned recording sites, and became larger moving toward more lateral sites. A significant placement versus cue-evoked dLight peak signal relationship was apparent early in training (R2=.545, p=.015), and became stronger at the late training phase (R2=.613, p=.0074). In contrast, in the dorsal striatum (DMS and DLS placements), early in training there was no relationship between cue-evoked dopamine signals and anatomical placement (R2=.104, p=.24). However, a significant relationship developed late in training (R2=.446, p=.0065), where the largest cue-evoked dopamine signals were measured in medially positioned placements, which became smaller as placements moved laterally. Within subjects, spontaneous cue responses in the early training phase were not significantly correlated with cue-evoked responses later in training (p=.943), indicating that they emerge as a function of learning. Together this data indicates that heterogeneous cue-encoding is driven by VTA dopamine neurons, with a dynamic spatiotemporal pattern across the striatum.

Given the specific recruitment of cue-evoked dopamine signals in the core and DMS, we focused additional analysis on the relationship between cue signals in these regions and the emergence of cue-conditioned behavior (**Fig 3**). We pooled trials in the early versus mid/late training phases across rats and plotted the peak cue-evoked dLight signal with trials sorted as a function of the average speed reached during the cue. Inspection of these heatmaps shows that cue-evoked signals were evident in the core early in training, but did not vary as a function of trial speed (**Fig 3A**, faster trials on top). In the later training stages, stronger and more enduring dLight signals following cue onset emerged on the faster movement trials (**Fig 3B**). In contrast, in the DMS, no consistent cue-evoked activity is seen early in training (**Fig 3C**), but strong cue signals emerge later, which don’t clearly differentiate movement speed (**Fig 3D**). We further analyzed this trial-by-trial data by running correlations of movement speed variables (max speed during cue, average speed during cue, and latency to max speed) against the peak cue-evoked dLight signal for each region. This revealed that NAc core cue-evoked signals were not correlated with movement vigor early in training (**Fig 3E**; max speed p=.901, avg speed p=.955, latency to max p=.739), but became consistently correlated at the mid (**Fig 3F**; max speed p=.001, avg speed p=.0002, latency to max p=.817) and late (**Fig 3G**; max speed p=.013, avg speed p=.0027, latency to max p=.039) training phases. In contrast, DMS cue-evoked signals did not clearly become correlated with movement variables across training (**Fig 3H-J**, all ps>.1), with the exception of a positive correlation with max speed in the late training phase (**Fig 3J**, max speed p=.0013). Collectively these results suggest that dopamine signals evoked by a VTA dopamine neuron predictive cue in the NAc core encode features of cue conditioned movement vigor earlier, and preferentially, compared to DMS signals.

**Figure 3.**
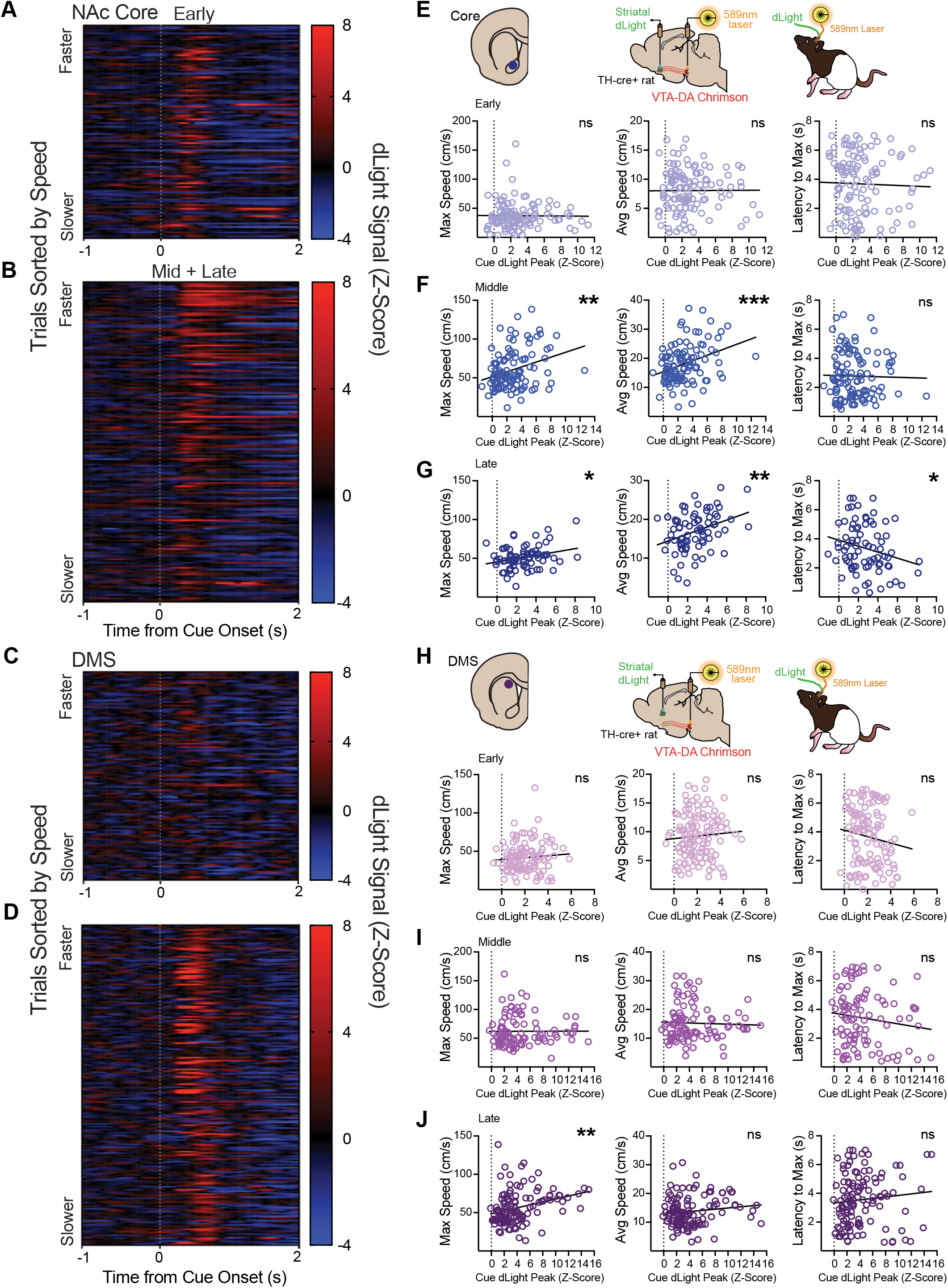
VTA cue-evoked dopamine signals in the NAc core preferentially predict the vigor of conditioned movement. A) Heatmap of Z-scored NAc core dLight signals for the early versus B) mid/late training phases centered around onset of cues predicting VTA dopamine neuron activation. Each row represents a trial, which are sorted as a function of the average speed reached during the entire cue for each trial, with faster movement trials on top and slower movement trials on the bottom. Corresponding heatmaps for C) early and D) mid/late DMS cue signals. E-G) Pearson correlations between the peak cue-evoked signal in the NAc core and the maximum speed, average speed, and latency to maximum speed, trial by trial, for each training phase. Cue-evoked dLight in the core was not correlated with any speed measure E) early in training, but positive correlations with maximum and average speed emerged in the F) middle and G) late training phases, while a negative correlation with latency to maximum speed emerged in the late training phase. H-J) Corresponding cue signal versus speed variable correlations for the DMS. Across these analyses, cue-evoked dLight in the DMS was only correlated with the maximum speed in the J) late training phase. ***p<.001, **p<.01, *p<.05.

### VTA dopamine neuron activation evokes DA signaling preferentially in medial striatum

Optogenetic activation of VTA dopamine neurons alters neural activity in a number of downstream regions, including throughout the striatum and parts of the frontal cortex, in rodents (52,53), but the impact on broad striatal dopamine signaling remains unclear. We next explored how VTA dopamine neuron activity alters dopamine signaling across the striatum (**Fig 4**). On dLight recording sessions, cues were paired with laser stimulation (20 Hz, 5 sec) on 80% of trials. Activation of VTA dopamine neurons during optogenetic conditioning produced a robust dopamine increase in the NAc medial shell, core, and DMS, and small responses in the DLS that were time-locked to laser delivery (**Fig 4B-E**). Focusing our analysis on the laser stimulation window, we found broadly heterogeneous dopamine signals across striatal recording sites. We quantified the average Z-score and AUC measures, which varied by region and training phase (**Fig 4F,G**, mixed effects ANOVA interaction of region and phase, F(6,38)=17.9, p<.0001); main effect of region, F(3,21)=9.11, p=.0005). In the medial shell, dopamine signals were overall largest, and increased in magnitude across training (**Fig 4B,F,G**; **Fig 4H**, paired t test for shell, t(3)=3.01, p=.029), which was not the case for other regions. In the core (**Fig 4C,F,G**; **Fig 3H**, paired t test for core, t(5)=.939, p=.196) and DMS (**Fig 4D,F,G; Fig 4H**, paired t test for DMS, t(5)=.762, p=.24), laser-evoked dopamine signals were stable across training phases. In the DLS, we saw little laser-evoked dopamine, and this small signal actually decreased with extended training (**Fig 4E,F,G**; **Fig 4H**, paired t test for DLS, t(8)=2.35, p=.047), despite the development of vigorous conditioned behavior. Given the large laser-evoked signal in the shell relative to the other regions, we separately analyzed signals in the core, DMS, and DLS, with the shell excluded. Stimulation-evoked dopamine remained variable across region and training phase (Interaction of phase and region, F(4,33)=4.15, p=.0078; main effect of region, F(2,18)=11.53, p=.0006). We next examined the relationship between VTA dopamine neuron stimulation and evoked dopamine signals across the sampled striatal space of our recording sites, independent of subgrouping. We found an overall gradient of laser-evoked dopamine, with the signal magnitude shrinking along the ventromedial (largest) to dorsolateral (smallest) axis in the coronal plane. Early in training, there was a trend for a correlation between recording placement in the striatum and dopamine signal (**Supplemental Fig 7**, R2=.14, p=.064), which became highly significant by the late training phase (R2=.41, p=.0005). Within subjects, across the whole dataset, stimulation-evoked dLight early in training was highly correlated with the signal evoked by laser later in training (p=.0003), indicating general stability in the activation of dopamine neurons throughout the experiment. Finally, we looked at the potential relationship between variability in stimulation fiber placement, which were clustered over center of the VTA (mean fiber location +0.69mm lateral) with a medial to lateral spread of about 600 microns (**Fig 1B**) and found no correlation early (p=.411) or late (p=.216) in training.

**Figure 4.**
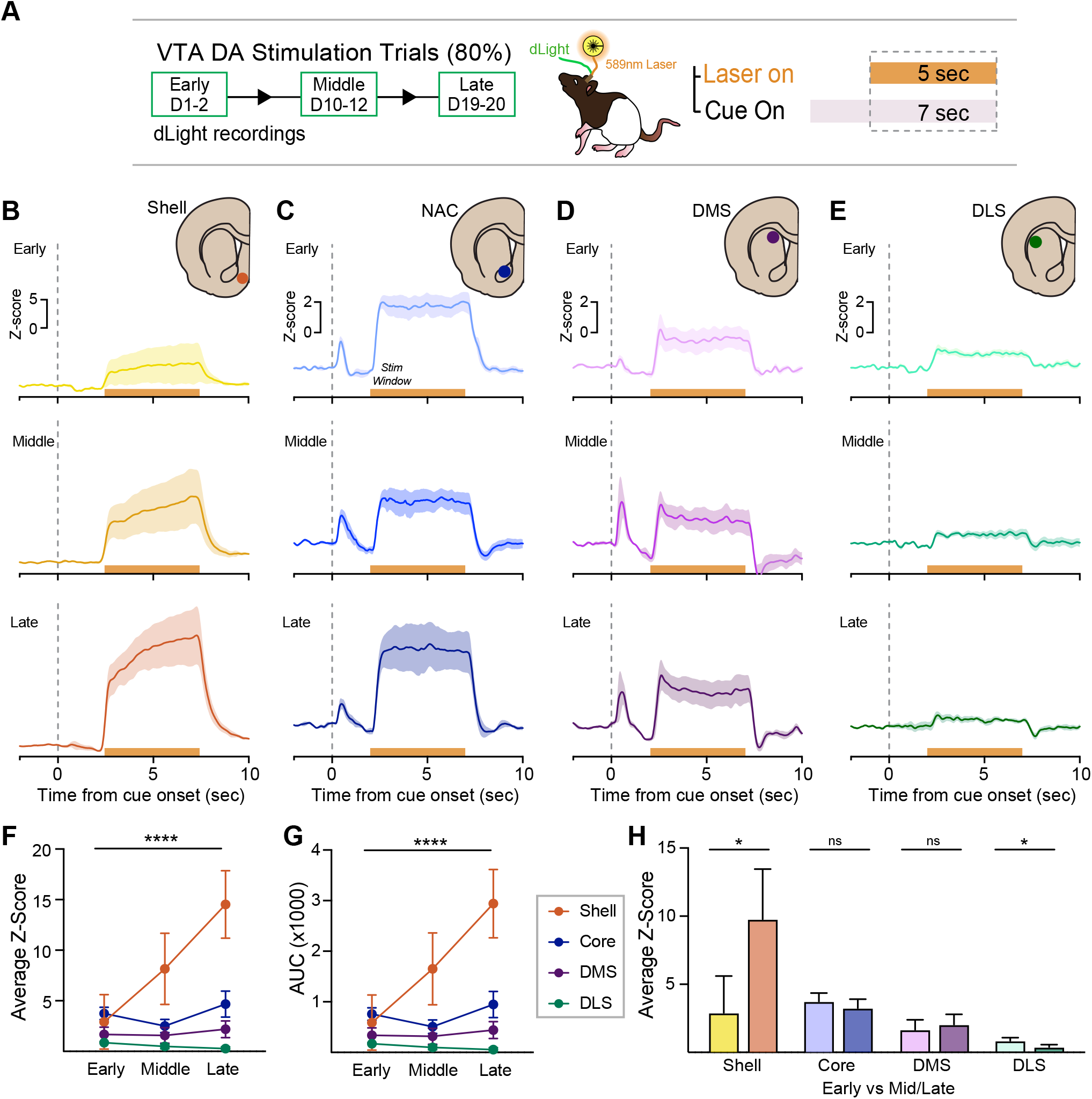
VTA dopamine neurons drive dopamine signaling preferentially in the medial striatum. A) Schematic of optogenetic Pavlovian conditioning paradigm. On dLight recording sessions, cues were paired with optogenetic stimulation of VTA dopamine neurons on 80% (20/25) of trials. Data in this figure are from the stimulation window, which corresponds to the final 5 sec of each cue period. B-C) Optogenetic activation of VTA dopamine neurons (20Hz, 5sec) produced different patterns of dopamine signaling across striatal targets. B) Large laser-evoked dopamine signals were present from training onset in the NAc medial shell, and this laser-evoked signal increased in magnitude across training. C) Laser evoked robust dopamine signals in the NAc core, which were stable across training. D) Laser also evoked robust dopamine signals in the DMS, which were stable across training. E) In contrast, optogenetic stimulation of VTA dopamine neurons evoked minimal dopamine signals in the DLS. F) Average dLight signal Z-score and G) AUC values varied significantly by region and training phase. H) Stimulation-evoked dLight signals increased in the shell, remained stable in the core and DMS, and decreased in the DLS, across training. ****p<.0001, *p<.05.

### Dopamine signals in the NAc core and DMS, but not shell or DLS, report dopamine prediction errors

A central feature of dopamine neuron activity is their ability to signal prediction errors, or the discrepancy between predicted and actual outcomes (12,13,54). We assessed whether striatal dopamine signals similarly report on a violation in the expectation of dopamine neuron activation (**Fig 5**). On dLight recordings sessions during optogenetic Pavlovian conditioning, 20% of trials were expectation probes, where the cue was presented normally, but laser stimulation was omitted (**Fig 5A**). Rats behaved similarly on omission trials, compared to laser stimulation trials, moving in response to cues for the same percentage of time (**Fig 5B**, mixed effects ANOVA no main effect of trial type, F(1,19)=1.81, p=.195), and reaching the same maximum speed across training phases (no main effect of trial type, F(1,19)=.748, p=.398). This further underscores that behavior in our optogenetic conditioning tasks reflected learned conditioned responses rather than simple laser-evoked movements. On omission trials, movement was tightly locked to cue-presentations (**Fig 5C**) and increased in speed and vigor across training phases (**Fig 5D-F**; average speed, F(1.70,27.11)=23.8, p<.0001; maximum speed, F(1.54,24.61)=19.04, p<.0001; time to maximum speed, F(1.48,23.72)=61.6, p<.0001), similar to stimulation trials (**Fig 1**).

**Figure 5.**
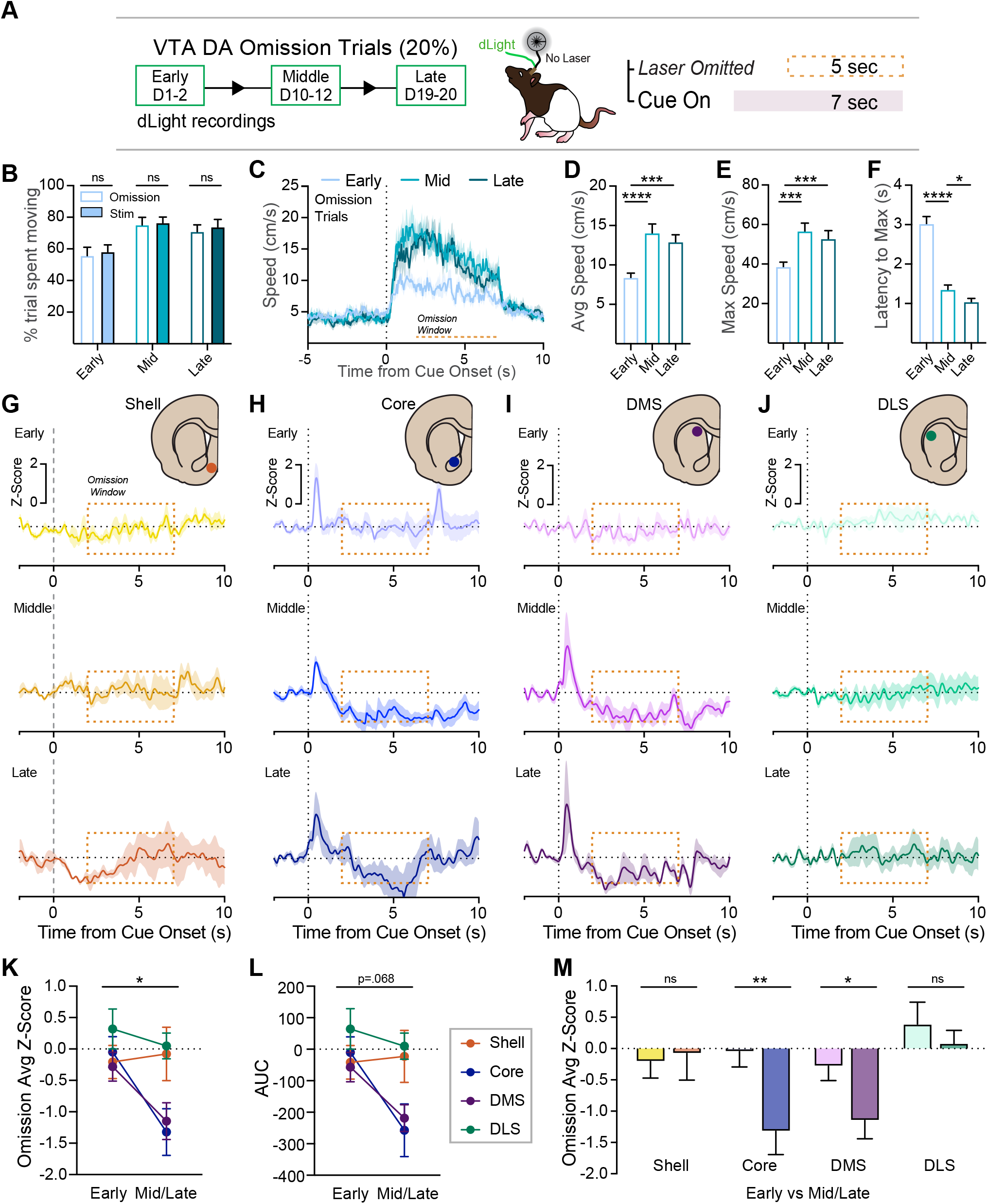
Dopamine signals in the NAc core and DMS, but not shell or DLS, report dopamine prediction errors. A) Schematic of optogenetic Pavlovian conditioning paradigm. On dLight recording sessions, laser delivery was omitted on 25% of cue presentations. B) Rats spent similar amounts of time moving during cue presentations on stimulation and omission trials. C) On omission trials, movement was tightly locked to cue onset and offset, increasing in speed across the early (days 1-2), middle (days 10-12), and late (days 19-20) phases of training. Both average D) and maximum E) speed achieved increased between early and middle/late training phases. F) Movement acceleration, reflected in the decreased time required to reach maximum speed, increased between each training phase. G-J) Omission of cue-paired optogenetic activation of VTA dopamine neurons led to heterogeneous dopamine responses across recording sites. G) Dopamine signals in the NAc medial shell did not systematically change during the laser omission window. H) In the NAc core, a laser omission-related dip in dopamine signals emerged across the middle and late training phases. I) Similarly in DMS, omission of VTA dopamine neuron stimulation resulted in a dip in dopamine at the middle and late training phases. J) Laser omission-related changes in dopamine signal did not emerge in the DLS at any training stage. K,L) During the omission window the average signal z-score and AUC varied across region and training phase. M) Omission-related signals decreased in the core and the DMS, but not in the shell and DLS across training. ****p<.0001, ***p<.001, **p<.01, *p<.05.

We found heterogeneous signals across recording sites during these laser-omission probes (**Fig 5G-J**). Dopamine signals during the laser omission window varied as a function of region and training phase (**Fig 5K**, average Z score, mixed effects ANOVA interaction of phase and region, F(3,21)=3.129, p=0.04; **Fig 5L**, AUC, trend for interaction of phase and region, F(3,21)=2.76, p=.068, main effect of region, F(3,21)=3.34, p=.039). During the window of omission of expected dopamine neuron excitation, dopamine signals in the NAc core and DMS dipped below baseline (**Fig 5H,I**), consistent with a negative prediction error. These core and DMS error signals were not present at the beginning of training, but developed in the later phases (**Fig 5M**, paired t test for core, t(5)=4.03, p=.005; paired t test for DMS, t(5)=2.31, p=.035). This echoes dopamine response patterns when natural rewards are omitted (49,54,55). In contrast, dopamine signals in the shell and DLS were unresponsive to laser omission, regardless of training phase (**Fig 5G,J,M**). Taking our cue-response data (**Fig 2**) and omission data (**Fig 5**) together, we find that only the NAc core and DMS engage in prediction error-like dopamine signaling based on learning driven by VTA dopamine neuron activity.

### Pavlovian learning driven by SNC dopamine neuron stimulation directs unique, heterogeneous striatal dopamine signals

So far, our results show that learning driven by VTA dopamine neurons does not recruit DLS dopamine signals. Critically, stimulation of SNC dopamine neurons is also sufficient to drive Pavlovian learning, but the content and behavioral impact of that learning is distinct from that engaged by VTA dopamine neurons (6,10,56). To build on this notion, we next explored the striatal dopamine impact of optogenetic conditioning of SNC dopamine neurons.

In a new cohort of TH-cre rats, cre-dependent Chrimson virus and stimulation optic fibers were targeted to SNC dopamine neurons (**Supplemental Fig 2**, mean fiber location +2.2mm lateral) and optogenetic Pavlovian conditioning was completed as above (**Supplemental Fig 8A-C**). Similar to the VTA conditioning groups, we found that cues predicting SNC dopamine neuron activation drove robust learning, as quantified by an increase in the likelihood of movement and movement speed during cue presentations (**Supplemental Fig 8D**). The average and maximum speed of conditioned movement during cue presentations increased across training (**Supplemental Fig 8E,F**; paired t-tests early vs late: t(9)=3.75, p=.005, t(9)=3.21, p=.011). As training progressed, we saw a general reduction in cue-directed approach behavior (**Supplemental Fig 8G**, % cue periods with an approach: mixed effects ANOVA main effect of training phase, F(1.88,11.25)=4.15, p = 0.046), and an increase in rotational behavior (**Supplemental Fig 8H,I**, angular speed: mixed effects ANOVA main effect of training phase, F(1.46,11.42)=11.42, p = 0.0046), in SNC conditioned rats, consistent with previous studies (6). As with VTA rats, SNC conditioned turning behavior was cue-evoked, and always occurred contralateral to the stimulation site.

During this SNC dopamine optogenetic conditioning (**Fig 6A**), we recorded dLight dopamine signals in the striatum, similar to above. We chose to focus specifically on the NAc core (n=9) and the DLS (n=5), as these regions showed the most distinct patterns in the VTA conditioning groups. Furthermore, while there is some evidence that SNC dopamine neurons may communicate broadly across the medial-lateral extent of the dorsal striatum to engage DLS and DMS specific activity patterns (57,58), it remains unknown whether SNC dopamine neurons engage dopamine signaling in the ventral striatum during learning.

**Figure 6.**
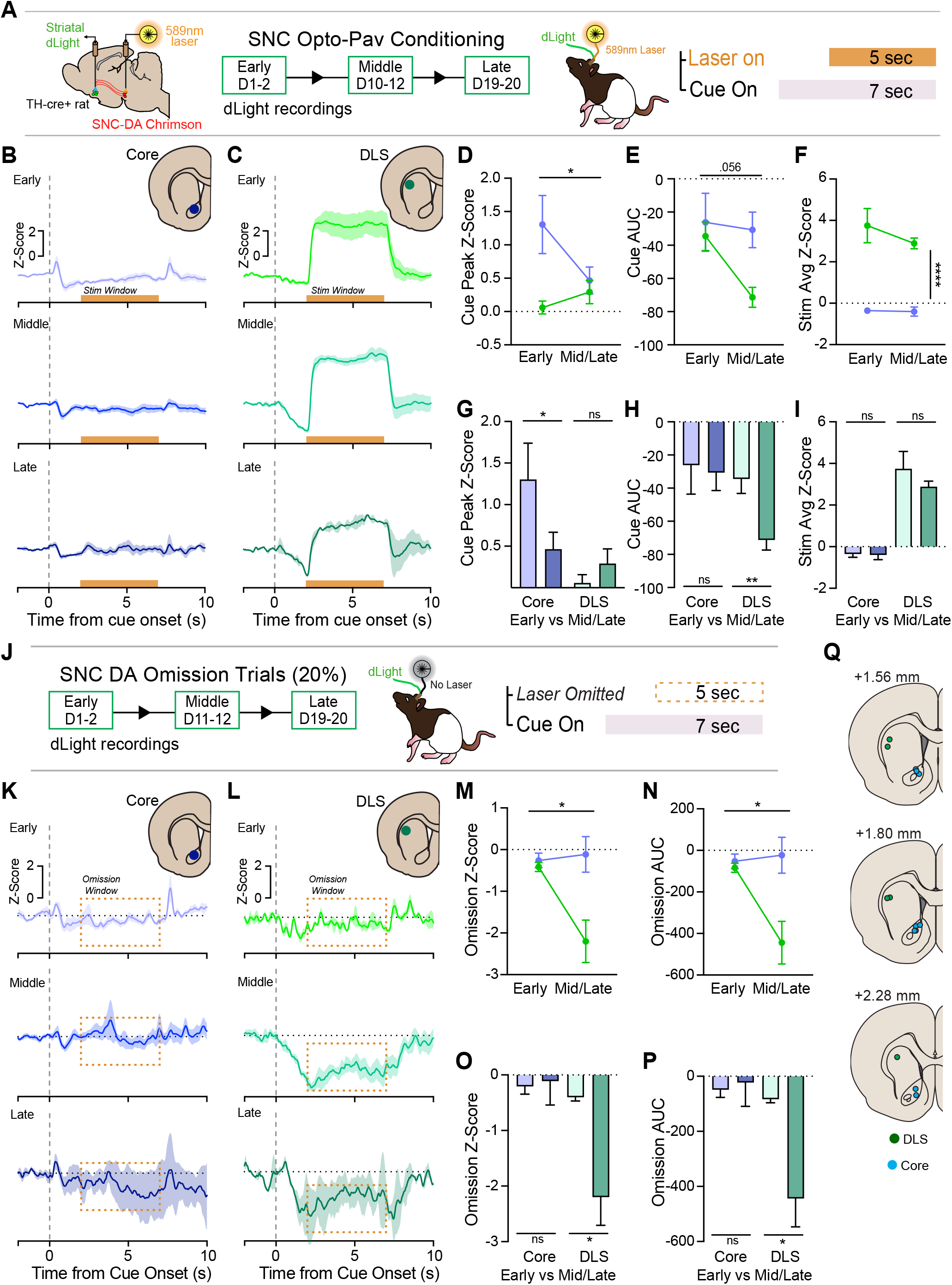
Heterogeneous striatal dopamine signals emerge during learning driven by SNC dopamine neuron stimulation. A) Overview of recording scheme during behavioral testing. Chrimson was targeted to SNC dopamine neurons and dLight signals were recorded in the NAc core (n=9) and DLS (n=5) across optogenetic Pavlovian conditioning. B,C) We saw heterogeneous dopamine signals in the core and DLS in response to cues predict ing SNC dopamine neuron stimulation (first 2-sec of cue), and the stimulation period itself. B) In the core, there was an initial cue response, which diminished in peak with training. C) In the DLS, no positive peak occurred, but a steady negative dip in dLight emerged in response to the cue across training. D,E,G,H) Quantification of the cue peak and AUC measures showed that the positive core peak diminished with training, and the cue AUC for the DLS became more negative with training. F,I) Strong laser-evoked dLight fluorescence was seen in the DLS across training, but not in the core at any training stage. J) Schematic of optogenetic Pavlovian conditioning paradigm for omission probes. On dLight recording sessions, laser delivery was omitted on 20% of cue presentations. K,L) Divergent dLight signals were seen in the core and DLS when laser stimulation of SNC dopamine neurons was omitted on a subset of trials. K) Core signals did not systematically change during omission. L) In the DLS, dLight fluorescence dipped at cue onset as training progressed. This signal maintained a negative deflection during omission trials, with a slight upward trend. Quantification of the M,O) z-score and N,P) AUC fluorescence during the omission period confirmed that core signals did not change with omission, but DLS signals became more negative across training. Q) Summary of recording fiber placements in the DLS and core. ****p<.0001, **p<.01, *p<.05.

Spatially and temporally heterogeneous dopamine signals emerged in response to SNC dopamine neuron predictive cues (**Fig 6B-I**). Brief positive cue-evoked signals were seen in the NAc core early in training and diminished as conditioning progressed (**Fig 6B**). In the DLS, no cue-evoked response was seen early, but across training, a slow negative dip following cue onset emerged (**Fig 6C**). We quantified the cue-evoked signal, which varied systematically in peak and AUC (**Fig 6D,G,E,I**; peak: region by phase interaction, F(2,12)=5.44, p=.038; AUC: trend main effect of phase, F(1,12)=4.44, p=.056). The peak signal in the NAc core diminished, while the DLS peak remained near zero (**Fig 6G**, paired t test for core, t(8)=2.55, p=.034; paired t test for DLS, t(4)=1.72, p=.16). The overall AUC of the cue-evoked dopamine signal was slightly negative in the core, and stable across training (**Fig 6H**, paired t test for core, t(8)=.329 p=.75). In contrast, the DLS AUC and trough/minimum response increased with training (**Fig 6H**, paired t test for DLS, t(4)=5.04, p=.007; paired t test for DLS minimum cue Z-score early vs mid/late, t(4)=6.22, p=.003). We examined the relationship between SNC cue-related signals and behavior on a trial-by-trial basis (**Supplemental Fig 9**). While the initial NAc cue peak signals were positively correlated with movement speed (p=.0001), that relationship went away as training progressed (**Supplemental Fig 9A,B,E,F**). Cue-evoked dips in dopamine in the DLS were not correlated with movement variables at any stage of training (**Supplemental Fig 9C,D,G,H**; ps>.2).

During the stimulation window, optogenetic activation of SNC dopamine neurons produced variable dopamine responses across recording sites (**Fig 6B,C,F,I**; average Z-score main effect of region: F(1,12)=140.4, p<.0001). In the core, dLight signal remained near zero during simulation, across phases (**Fig 6F,I**; paired t test for core, t(8)=.162, p=.86). In the DLS, strong laser-evoked dopamine signals were consistent across each phase (**Fig 5F,I**; paired t test for DLS, t(4)=.89, p=.42). For further comparison of learning-related dopamine signals in the NAc core and DLS, we plotted data for VTA and SNC Pavlovian conditioning groups together on the same graphs (**Supplemental Fig 10**). This analysis shows that learning driven by VTA versus SNC dopamine neurons, despite promoting strong conditioned behavioral invigoration in either case, result in nearly diametrically opposed dopamine signals in the dorsal and ventral striatum.

We next examined dopamine signals during laser omission trials (**Fig 6J**), finding different patterns in the core and DLS (**Fig 6K,L**). Quantification of these signals revealed that average signal Z-score and AUC during the omission window varied across regions and phases (**Fig 6M-P**; average Z-score region by phase interaction, F(1,20)=4.80, p=.039; main effect of region, F(1,20)=6.49, p=.019; AUC region by phase interaction, F(1,20)=4.84, p=.04). In the NAc core, omission of SNC dopamine neuron stimulation during conditioning had no effect on dopamine signals, which remained at zero/baseline (**Fig 6O,P**; average Z-score paired t test for core, t(8)=.209, p=.84; AUC paired t test for core, t(8)=.28, p=.79). In contrast, negative DLS dopamine signals during the omission window emerged across training (**Fig 6O,P**; average Z-score paired t test for DLS, t(4)=3.38, p=.028; AUC paired t test for DLS, t(4)=3.39, p=.0275). Based on qualitative inspection of the DLS omission traces (**Fig 6L**), however, the increasingly negative Z-score and AUC values during the omission window appear to reflect that the DLS signal was already negative following cue onset, an effect that grew across conditioning. This suggests that omission related DLS signals following SNC conditioning are potentially unique from those seen in the core and DMS during VTA conditioning (**Fig 5**), and they do not necessarily indicate a negative prediction error.

## DISCUSSION

Here, we characterized the emergence of cue-evoked behavior as a consequence of learning driven by dopamine neuron activation. Recording dopamine signals with fiber photometry across different striatal regions during optogenetic Pavlovian conditioning, we show that there is considerable heterogeneity in the extent that VTA and SNC dopamine neurons can drive learning-related dopamine signals downstream. In the ventral striatum, we found that cues predictive of VTA dopamine neuron activation evoked dopamine signals in the NAc core. In contrast, cue-evoked signals did not emerge in the nearby NAc medial shell, despite VTA dopamine neuron excitation producing a robust increase in dopamine there, In the dorsal striatum, cue and laser-evoked dopamine signals also emerged in the DMS, with cue-evoked signals developing after NAc core signals. Surprisingly, extended learning mediated by VTA dopamine neuron activity was not sufficient to engage robust cue or laser-evoked dopamine signals in the DLS, despite promoting rapid conditioned movement and stereotyped behavior patterns. Further, we found that NAc core and DMS, but not NAc shell or DLS dopamine signals, report prediction errors by dipping in magnitude when expected dopamine neuron stimulation was omitted. Finally, learning driven by SNC dopamine neuron stimulation failed to engage clear cue or expectation-related dopamine signals, despite promoting vigorous movement. Taken together, these results indicate dopamine neurons engage considerable spatial and temporal complexity in the striatum during learning. These results offer new insight into heterogeneity in striatal dopamine function, as well as the in vivo circuit architecture of the midbrain-striatum network.

### Specialized cue, error, and reward signals across striatal dopamine targets

Dopamine signals in response to cues predicting optogenetic activation of VTA dopamine neurons developed specifically in the NAc core and the DMS, but on different timescales. Core signals were unique among all recorded areas, as they were apparent at the beginning of conditioning, when cue-evoked movement patterns were less vigorous and more directed at the cue. Our data suggest the function of the NAc core signal evolves over time, where early signals reflect some aspect of novelty or general salience (46) and later signals reflect motivational vigor. Cue-evoked signals in the DMS emerged later in training, roughly in parallel to the stabilization of more rapid movement speeds and reduction in approach behavior. We also found error-related signaling in the NAc core and DMS, where dopamine signals dipped below baseline at the exact time when dopamine stimulation was omitted. In both regions, error signals only emerged after learning had occurred. The fact that this type of striatal plasticity can be engaged in the absence of learning driven by interface with a natural reward suggests that it is a fundamental feature of the system. The lack of error-related DLS dopamine fits with evidence that dopamine neuron axons in the DLS are less responsive to negative prediction errors compared to other regions (30).

Based on our data, we conclude that NAc core and DMS dopamine are unique among striatal subregions in that they prioritize encoding of expectation-related information. The broad similarity we see in NAc core and DMS dopamine signals during Pavlovian learning is interesting in light of a number of studies that show DMS and core dopamine signals diverge and have unique functional roles during instrumental decision making tasks (29,59,60). One possibility is that while nominally similar to the core, the DMS cue signals in our studies could be more related to action values and action prediction, which reflect learned cues and decision states that function somewhat independent of reward value or reinforcement (59,61–63). The fact that we saw a tighter relationship between cue-evoked dopamine and behavioral invigoration in the core versus the DMS supports a framework where core signals preferentially encode the motivational features of Pavlovian cues (42,64,65). In other studies, major differences for the role of VTA and SNC dopamine neurons in reinforcement, cue valuation, and model-based learning are becoming clear (6,7,10,56), but direct comparisons between DMS and NAc core projecting dopamine neurons in these frameworks remain mostly unexplored.

Notably, we found robust VTA laser-evoked dopamine signals in the DMS, in addition to the NAc core and shell. This suggests that dopamine neurons projecting to the NAc and DMS are somewhat intermingled in the VTA/medial SNC, allowing for an interface between dopamine in the medial portion of the striatum across the dorsal-ventral axis. This idea is also supported by anatomical work (14,15), but it will be important to compare our results with future studies where VTA dopamine neurons that project only to the NAc are targeted. Previous studies using functional imaging to measure brain activation generated by dopamine neuron stimulation support the notion that dopamine neurons have broad access to alter network level activity (52,53). Given this, the lack of clear dopamine signals in the DLS following VTA dopamine neuron activation is perhaps surprising. It remains possible that intrinsic DLS neurons were engaged during cue or laser delivery, but it is notable that we actually saw a decrease in the small DLS dopamine signal generated by VTA activation. Another possibility is that DLS dopamine signals changed in some general way, potentially measurable outside of cue-trial periods on which we focused. While we did not see an change in baseline movement speed outside of cue and laser periods across conditioning, VTA dopamine based learning could have resulted in subtle changes in the organization of reward-independent spontaneous behavior, which is thought to engage DLS dopamine (66). Thus, DLS changes may have occurred, but not in an obvious gain-of-function capacity. Further, it will be critical to assess whether neural activity in DLS or DMS is required for the emergence of vigorous cue-evoked movement during dopamine-mediated learning.

In the NAc medial shell, we found a wholly unique dopamine pattern compared to the other recording sites. There was no evidence of either cue or error-related signals in the shell, but instead, the largest laser-evoked dopamine signals. This motivates an important conclusion borne out by several features of our data: the degree to which a striatal site can be conditioned to encode cue-related information is not simply a matter of how much dopamine is stimulated into it. Recent evidence indicates anatomically localized functions for the shell may be due to unique terminal mechanisms (67), as well as the specific patterns of connectivity with the midbrain (33,68,69), compared to other parts of the ventral striatum. Notably, our data reflect dopamine recordings in the dorsal portion of medial shell, and other recent work indicates that conditioned stimuli evoke unique encoding mechanisms along a dorsoventral axis within the medial shell (33,70,71), which require further investigation. Overall our results fit with the general notion that dopamine in the medial shell preferentially signals reward and reward value over other features of learning (72–75). They also support anatomical frameworks classifying the medial shell as part of a distinct brain macrosystem within limbic networks (76,77).

The results from our SNC conditioning experiments motivate a number of additional considerations. First, we find that optogenetic stimulation of SNC dopamine neurons does not evoke dopamine release in the NAc core, and SNC-predictive cues do not drive conditioned dopamine responses or prediction errors in the core. These results give an in vivo explanation for why, across a slew of studies, and in classic conceptual frameworks of dopamine heterogeneity, SNC dopamine manipulations are shown to support fundamentally different learning processes (2,4,6,10,56,61,64,78). SNC dopamine neurons are generally unable to imbue cues and actions with conditioned value to promote flexible reward seeking, a function that seems to rely on or require activity in VTA dopamine neurons projecting to the ventral striatum. The presence of novelty cue responses in the core early in SNC dopamine conditioning in our data, which diminish as training progresses, lends further support to the notion that early responses in the core are qualitatively different from those emerging later in training in the VTA conditioning groups. Critically, our results show that at least some forms of Pavlovian learning do not require the development and maintenance of cue-evoked dopamine signatures in the NAc core.

We find unique cue-evoked signals in the DLS in response to learning driven by SNC dopamine neurons. Despite strong laser-evoked dopamine release in the DLS and robust conditioned behavior, dopamine responses there to the SNC-predictive cue were consistently negative, a trend that increased across training. Based on the current results alone these conditioned dips are difficult to fully explain. Phasic activation of SNC dopamine neurons is strongly reinforcing (6,10,56,79), and fluctuations in DLS dopamine are thought to facilitate exploration and shape movement sequences (8,66,80), so it is unlikely that the negative cue signals seen here reflect conditioned aversion. Instead, this DLS dopamine signature may stem from the unique striatal network engagement that SNC dopamine neurons promote, compared to VTA dopamine neurons. SNC dopamine neurons may recruit other circuits, or terminal mechanisms controlling dopamine release locally, that briefly suppress DLS dopamine signaling during learning. Results from SNC cell body recording studies have shown mixed results, with examples of both increases and decreases in dopamine cell firing before movement initiation (27,81,82), and subtypes of SNC dopamine neurons respond to movement and reward differently (38). The connection to striatal dopamine release in our studies from those results is unclear, but our data indicate that through SNC-mediated learning, dopamine dips are engaged at the level of the striatum. In other studies, fluctuations in cholinergic interneurons gate dopamine release in the dorsal striatum via extrastriatal glutamatergic inputs (83–85), and so it will be important to investigate how dopamine neuron mediated learning engages other components of the corticothalamostriatal network. Future studies will be needed to parse these factors, but regardless of the specific mechanism, our results support a framework where VTA and SNC dopamine neurons orchestrate qualitatively different learning mechanisms, via unique engagement of striatal dopamine.

### Implications for the functional architecture of the midbrain-striatum network

The current study is unique in that we isolate the impact of learning driven directly by dopamine neurons and allow determination of how dopamine neurons engage the broader striatal network, for insight on long-standing questions about its functional organization. In the classic “ascending” spiral framework (16,17), behavioral control and neural signals transition from VTA inputs to the ventral striatum to SNC inputs to the dorsal striatum (18–20) in parallel with stereotyped, habitual action patterns seen after extended learning (21). Our results potentially offer some support for this, given that conditioned dopamine signaling engaged by VTA dopamine neuron mediated learning first appeared in the NAc, then later emerged in the DMS, when conditioned behavior was more vigorous. Critically, we show that rather than a shift away from ventral striatum, an additive, progressive recruitment of DMS occurs instead. This recruitment also appears to be limited to medial dorsal regions, as we did not see DLS dopamine activity emerge, even after extensive training.

Overall, our work fits with recent ex vivo work demonstrating that DMS-projecting dopamine neurons do not have circuit-level functional connectivity with DLS-projecting dopamine neurons (58). Rather, midbrain-striatal systems exist as reciprocal, parallel, primarily independent loops, where VTA dopamine neurons can control their own (but not SNC) activity via direct pathway projections from the ventral striatum (17,57,58,68,86). This suggests that at least in naive mice, there is no latent connectivity to support an ascending DLS engagement, but a critical component of the spiral framework is that learning drives plasticity that recruits a spiral mechanism. Our work explores this notion in vivo, and we find little direct evidence: despite extended Pavlovian learning via VTA dopamine neuron activation that produces rapid cue-evoked movement patterns, neither cue nor robust laser-evoked dopamine signals emerge in the DLS. Further, NAc core dopamine signals actually became more correlated with conditioned movement invigoration across training. Consistent with this, NAc, but not DLS dopamine signaling and neural activity remain important for normal execution of Pavlovian conditioned cue approach after extended training (28). The type of learning framework is likely key for how striatal networks are engaged, but this is not restricted to Pavlovian learning, as other recent work highlights a lack of a shift toward DLS dopamine signals and striatal activity during the development of outcome-insensitive instrumental habits (24–26). Our SNC conditioning experiments further indicate that “descending” recruitment of ventral striatum is not engaged by learning driven by dorsal striatum dopamine inputs. Collectively, our data support an updated account of the functional architecture of midbrain-striatal loops, where learning is not predicated on a consistent shift along the ventromedial to dorsolateral axis, or vice versa, in dopamine signaling.

It is important to consider our current results in the context of seminal studies demonstrating a clear shift in striatal recruitment to DLS (19,20,87,88). The strongest evidence for this effect comes from instrumental conditioning studies where cocaine is self administered for extended periods. This suggests that chronic hyperactivation of the dopamine system via drug stimuli may be necessary to promote DLS recruitment. Actual commerce with an external reward is not essential, however, as dopamine neuron self stimulation can promote plasticity in the dorsal striatum that is similar to that produced by cocaine self administration, independent of other reward exposure (89,90). Thus, DLS recruitment may depend on whether or not mesolimbic dopamine circuits are activated in a rapid, binge-like, or escalating fashion. Another possibility in our studies is that the DLS was conditionally recruited via mechanisms that are not reported by changes in dopamine release, including potentiation of corticostriatal inputs (91,92). Recent efforts have also characterized functionally distinct roles for striatal output pathways in behavioral control, via parallel channels through the thalamus, midbrain, and cortex (86,93,94). It will be important to investigate how dopamine neuron activity may engage such broader nigro-thalamo-cortical and striatal loops, which could provide a circuit mechanism by which midbrain inputs to the ventral striatum can engage dorsal striatum, independent of the classic spiral mechanisms. The recent discovery of pan-striatal wave-like activation patterns among dopamine signals and cholinergic interneurons provides another possible source of ventral-dorsal interactions (95,96).

## Conclusions

Here, we uncover fundamental region-specific patterns of dopamine signaling engaged during Pavlovian learning by measuring dopamine signals and behavior emergent from learning driven by dopamine neurons in the absence of other rewards. Our results show that VTA and SNC dopamine neurons direct learning about environmental cues by driving broad plasticity in the dopamine system, where preferential reward, cue, and error-related signaling in subregions of the ventral and dorsal striatum develop across learning phases. This work provides new insight into the functional architecture of the striatal network, building on long-standing anatomical and in vivo recording frameworks positing a shift in striatal control to the dorsal striatum as learning progresses. We find that dopamine neurons do not drive such a shift, despite the development of vigorous, cue-controlled movement patterns. Instead, as learning progresses, a broader landscaping of dopamine release patterns emerges across a network of ventral and dorsal striatal targets to uniquely encode movement, reward, and prediction-related features of learning.

## METHODS

### Subjects

Male and female TH-cre transgenic rats (n=30; 18F, 12M) bred on a Long-Evans background were used. These rats express Cre recombinase under the control of the tyrosine hydroxylase (TH) promoter (43). All rats weighed 250-500g at the time of surgery and were 4-7 months old at the time of experimentation. Experimental procedures were approved by the Institutional Animal Care and Use Committee at the University of Minnesota and were carried out in accordance with the guidelines on animal care and use of the National Institutes of Health of the United States.

### Surgical Procedures

Rats were anesthetized with 5% isoflurane and placed in a stereotaxic frame, after which anesthesia was maintained at 1-3%. Rats were administered saline, carprofen anesthetic (5 mg/kg), and cefazolin antibiotic (70 mg/kg) subcutaneously prior to start of surgery. The top of the skull was exposed and holes were made for viral infusion needles, optic fiber implants, and 4 skull screws. A total of 1.2 µL of virus (pAAV5-Syn-FLEX-Chrimson-tdTomato, Addgene) was infused unilaterally into the VTA (-5.6 AP, +/- 0.7 ML, -8.0 DV). In the same hemisphere, a virus coding for the dopamine biosensor dLight (39) (AAV5-hSyn-dLight1.3b, University of Minnesota Viral Vector and Cloning Core) was infused into the NAc (1.3 AP, +/- 1.3 ML, -7.0 DV), DMS (1.3 AP, +/1.3 ML, -4.3 DV), or DLS (1.7 AP, +/- 4.0 ML, -5.0 DV) at a total volume of 0.8 µL. A separate group of rats received the same Chrimson virus targeted to the SNC (-5.6 AP, +/- 2.5 ML, -7.5 DV) and dLight injections targeted to the ipsilateral DLS or NAc core. All viruses were infused at 0.1µL/min. Immediately after injection, the needle was raised 100 µm and then left in place for an additional 10 min to allow for diffusion. Optical fibers (9mm length, 400µm diameter, Doric Lenses) for photometry recordings were chronically implanted into the NAc (1.3 AP, +/- 1.3 ML, -6.6 DV), DMS (1.3 AP, +/- 1.3 ML, -4.3 DV), or dorsal striatum (1.7 AP, +/- 4.0 ML, -4.4 DV), and optic fibers (10mm length, 300µm diameter, Thorlabs) for optogenetic stimulation were implanted above the VTA (-5.6 AP, +/- 0.7 ML, -7.9 DV) or SNC (-5.6 AP, +/- 2.5 ML, -7.3 DV) in the same hemisphere as the viral infusions. All coordinates are in mm relative to bregma and skull surface. Implants were secured to the skull with dental acrylic (Lang Dental) applied around the skull screws and the base of the ferrules containing the optic fibers. At the end of all surgeries, topical anesthetic and antibiotic ointment were applied to the surgical site, rats were placed on a heating pad and monitored until they were ambulatory. After surgery rats were individually housed with ad libitum access to food and water on a 0800 to 2000 light/dark cycle (lights on at 0800), given carprofen (5 mg/kg) and cefazolin (70 mg/kg) for the first 3 days following surgery and weighed and monitored for 6 days. Optogenetic manipulations commenced at least 4 weeks after surgery. Post hoc validation of optic fiber placements revealed subsets of rats with fibers targeted to the core versus medial shell of the NAc, which we analyzed separately.

### Optogenetic Stimulation

Optogenetics studies used 589 nm lasers (OptoEngine and Dragon Lasers). Light output during individual 5-ms light pulses was adjusted to be approximately 2 mW/mm2 at the tip of the intracranial fiber (10-20mW constant power). For all optogenetic studies, optic tethers connecting rats to the rotary joint were sheathed in a lightweight armored jacket to prevent cable breakage and block visible light transmission. Laser pulses were initiated from Med-PC TTL pulses and patterned via Synapse software that also controlled photometry and video data acquisition.

### Fiber Photometry

To assess dopamine signaling across the striatum during Pavlovian conditioning, we measured dLight fluorescence in the NAc core, NAc shell, DMS, and DLS using fiber photometry. A fluorescence mini-cube (Doric-Lenses) transmitted light streams from 465 nm and 405 nm LEDs (Doric-Lenses), sinusoidally modulated at 211 Hz, and 330 Hz, respectively. LED power was set at ∼100 μW. On recording days rats were tethered to a core, shell, DMS, or DLS implant using a low autofluorescence cable sheathed in a lightweight armored jacket (Doric-Lenses). On all photometry recording days the rats were simultaneously tethered to the VTA or SNC implant. Photometry recordings were conducted on specific days with interspersed recording free days. Recording data here are sampled from the early (day 1 and 2), middle (days 10 12) and late (day 19 and 20) stages of Pavlovian conditioning.

Fluorescence from below the fiber tip was transmitted back to the mini-cube, where it was passed through a GFP emission filter, amplified, and focused onto a high sensitivity photoreceiver (Newport, Model 2151). A real-time signal processor (RZ5P, Tucker Davis Technologies) running Synapse software modulated the output of each LED and recorded photometry signals, which were sampled from the photodetector at 6.1 kHz. Demodulation of the brightness produced by the 465-nm excitation, which stimulates dopamine-dependent dLight fluorescence, versus isosbestic 405-nm excitation, which stimulates dLight in a dopamine-independent manner, allowed for correction for bleaching and movement artifacts. Task events (e.g., cue and laser presentations), were time stamped in the photometry data file via a TTL signal from the Med-PC behavioral program, and behavior was video recorded at 10-20 FPS, with timestamps for each frame captured in the photometry file.

### Habituation and Optogenetic Pavlovian Training

Optogenetic Pavlovian training was conducted as described previously (6). Briefly, rats were first acclimated to the behavioral chambers (Med Associates), conditioning cues, and optic cable tethering in a ∼30-min habituation session. During this session, rats were tethered to a rotary joint and 20 cue presentations, with no other consequences, were presented on a 90-s average variable time (VT) schedule. In each of the subsequent conditioning sessions, rats were presented with 25 cue (light + tone, 7 s) – laser stimulation (100 5-ms pulses at 20 Hz; laser train initiated 2 s after cue onset and continued for duration of cue) pairings delivered on a 100-sec VT schedule. The sessions lasted ∼42 min. These cues were never associated with another external stimulus (e.g., food or water). Cue and laser delivery were never contingent on an animal’s behavior and all rats received the same number of cue and laser events. On photometry recording days, 20% (5/25, pseudorandomly delivered) of trials served as dopamine omission probes, wherein cues were presented as normal, but laser stimulation did not occur.

### Video Scoring

During Pavlovian conditioning sessions behavior was video recorded using cameras (Vanxse CCTV Security Camera) positioned a standardized distance above each chamber. Videos from sessions 1, 2, 4, 8, 11, 12, and 19, 20 were scored offline by observers who were blind to the identity and anatomical target group of the rats. For days 1, 4, 8, 12, and 20, each cue-only (2-sec, 25 per session) and cue+laser (5-sec, 20 per session) event was scored for the occurrence and onset latency of the following behaviors. Locomotion: Defined as the rat moving all four feet in a forward direction (i.e., not simply lifting feet in place). Cue Approach: Defined as the rat’s nose coming within 2.5 cm of the cue light. Approach often involved the rat moving from another area of the chamber to come in physical contact with the cue light while it was illuminated. Rearing: Defined as the rat lifting its head and front feet off the chamber floor, either onto the side of the chamber, or into the air. Rotation: Defined as the rat making a complete 360-degree turn in one direction. For cue approach and rotation, the number of occurrences were also scored. For days 2, 11, and 19, only locomotion was scored.

### DeepLabCut pose estimation

Markerless tracking of animal body parts was conducted using version 2.2.1.1 of the DeepLabCut (DLC) Toolbox (40,44) and analysis of movement features based on these tracked coordinates was conducted in Matlab R2020b (Mathworks). All DLC analysis was conducted on a Dell G7-7590 laptop running Windows 10 with an Intel Core i7-9750H CPU, 2.60Ghz, 16 GB RAM, and an NVIDIA GeForce RTX 2080 Max-Q 8GB GPU. DeepLabCut 2.2.1.1 was installed in an Anaconda environment with Python 3.8.4, CUDA 11.7 and Tensorflow 2.10.

### DeepLabCut Model

2090 frames from 35 videos (32 different animals, 3 experiments) were labeled and 807 outlier frames were relabelled to refine the network. Labeled frames were split into a training set (95% of frames) and a test set (5% of frames). A ResNet-50 based neural network (45) was used for 1,030,000 training iterations. After the final refinement we found the test error was 4.1 pixels, the training error was 3.13 pixels and with a p-cutoff of 0.85 training error was 2.99 pixels and test error was 3.68 pixels. The body parts labeled included the nose, eyes, ears, center of head or fiber optic implant, shoulders, tail base, and an additional three points along the spine. Features of the environment were also labeled, including the 4 corners of the apparatus floor, two nose ports, two cue lights, two magazine ports, and 3 LED indicator lights when active (see **Supplemental Fig 1** for examples of labeled body parts and environment features).

### Behavior Analysis

DLC coordinates and confidence values for each bodypart and frame were imported to Matlab and filtered to exclude body parts/features from any frame where the confidence was < 0.7. For labeled features of the environment, which have a fixed location, the average coordinates for that recording were used for analysis. To convert pixel distances to the real chamber dimensions, for each video, a pixel to cm conversion rate was determined. The distance (in pixels) between each edge of the environment floor and the diagonal measurements from corner to corner were measured, and these values were divided by the actual distance in cm. The mean of these values was then used as the conversion factor. Movement speed was calculated from the implant coordinates frame by frame using the formula: [distance moved (pix per cm) * framerate] to give movement speed in cm/s. For plotting of speed data across recordings with different sampling rates, missing data points for the lower sampling rate sessions were filled in using linear interpolation with the interpol1 function (MATLAB), interpolated data points were not used to calculate peak or mean speeds for analysis. Approach to the cue light was detected by finding frames in which the nose was within 2.5 cm from the cue. The beginning and end of each approach occurred when the nose was further than 4.5 cm from the cue for 0.2s. Only approaches longer than 0.2 s were considered. Locomotion was detected using the speed of the 4 positions on the spine to detect whole-body movements. The movement threshold for detecting locomotion was calibrated to animal size using a scale factor determined from the relationship between body size (distance between the shoulder and mid-back points) and the optimal threshold for detecting movement in a separate group of animals not used for this study (n = 4). Locomotion bouts were detected when all of the visible body parts were above the movement threshold and a sliding window was used to determine when the speed of 2 or more body parts reached less than 50% of the detection threshold for 0.3s, indicating the beginning and end of a movement. Movements less than 0.5s in duration were excluded. Orientation toward the cue light was determined by calculating the angle between the vector from the implant to the nose with the vector from the implant to the cue. Animals were labeled as oriented to the cue whenever this angle was <30° for more than 0.2s.

### Photometry data collection and analysis

Photometry data was analyzed using a custom MATLAB pipeline, based on established methods for dLight data (24,39,46) and example code provided by TDT (Lick Bout Analysis.m). For analysis, signals were filtered with a 2-Hz lowpass filter, downsampled to 40 Hz, and a least-squares linear fit was applied to the 405-nm signal, to align it to the 465-nm signal. This fitted 405-nm signal was used to normalize the 465-nm signal, where ΔF/F = (465-nm signal – fitted 405-nm signal)/(fitted 405-nm signal). Normalized signals for each trial were extracted for 5s before and 10s after each cue presentation and Z-scored to the 5s pre-cue period for each trial to minimize the effects of drift in the signal across experiment duration. Cue responses were detected in the average trial waveform for each animal in the 2s window beginning at cue onset (and prior to optogenetic stimulation). Responses to the presence or absence of optogenetic stimulation were detected in the average trial waveforms for each animal across the 5s stimulation window, beginning 2s after cue onset. The maximum and minimum responses in each window were detected and latency was calculated relative to the start of the detection window. In some cases, the peak signal did not change, and so we also calculated waveform slopes, as a proxy for other features of dopamine signaling dynamics. The slope of the rising and falling phase of the peak and trough waveforms were calculated between the maximum and 50% of max using the formula t2-t1/f2-f1, where t is the beginning and end timepoints of the rising or falling phase of the waveform, and f is the Z-scored fluorescence at each timepoint. Area under the curve (AUC) values were calculated from the Z-scored traces by numerical integration via the trapezoidal method using the trapz function (Matlab). Peak and AUC of cue responses were calculated in the window after cue onset, before laser stimulation, to quantify the magnitude of cue-related signal changes. AUC for the stimulation window was calculated over the laser-on window. Recording groups were as follows: VTA-Dopamine conditioning: NAc shell (n=4), NAc core (n=6), DMS (n=6), and DLS (n=9); SNC-Dopamine conditioning: NAc core (n=9), DLS (n=5).

### Statistics

Behavior and photometry signal data were analyzed with a combination of linear mixed-effects ANOVA models, planned t tests, and Pearson correlations. Post-hoc comparisons were completed with Bonferroni corrections. Summary graphs represent averages of recordings sites/subjects, not individual trials. The data are expressed as the mean ± s.e.m. Statistical significance was set at p<0.05.

### Histology

Rats were deeply anesthetized with sodium pentobarbital (2 mL/kg) and transcardially perfused with cold phosphate buffered saline followed by 4% paraformaldehyde. Brains were removed and post-fixed in 4% paraformaldehyde for ∼24 hours, then cryoprotected in a 25% sucrose solution for at least 48 hours. Sections were cut at 50 microns on a cryostat (Leica CM1900). To confirm viral expression and optic fiber placements, brain sections containing the midbrain were mounted on microscope slides and coverslipped with Vectashield containing DAPI counterstain. Fluorescence from Chrimson and dLight as well as optic fiber damage location was then visualized using a Keyence BZ-X microscope.

## Acknowledgments

This work was supported by NIH grants F32 DA051138 (VC) and R00 DA042895, R01 MH129370, and R01 MH129320 (BTS). We thank Michael Dahl for assistance, and all members of the Saunders lab for their input and support in completion of this work.

The authors declare no competing interests.

## Supplemental Figures

**Supplemental Figure 1.**
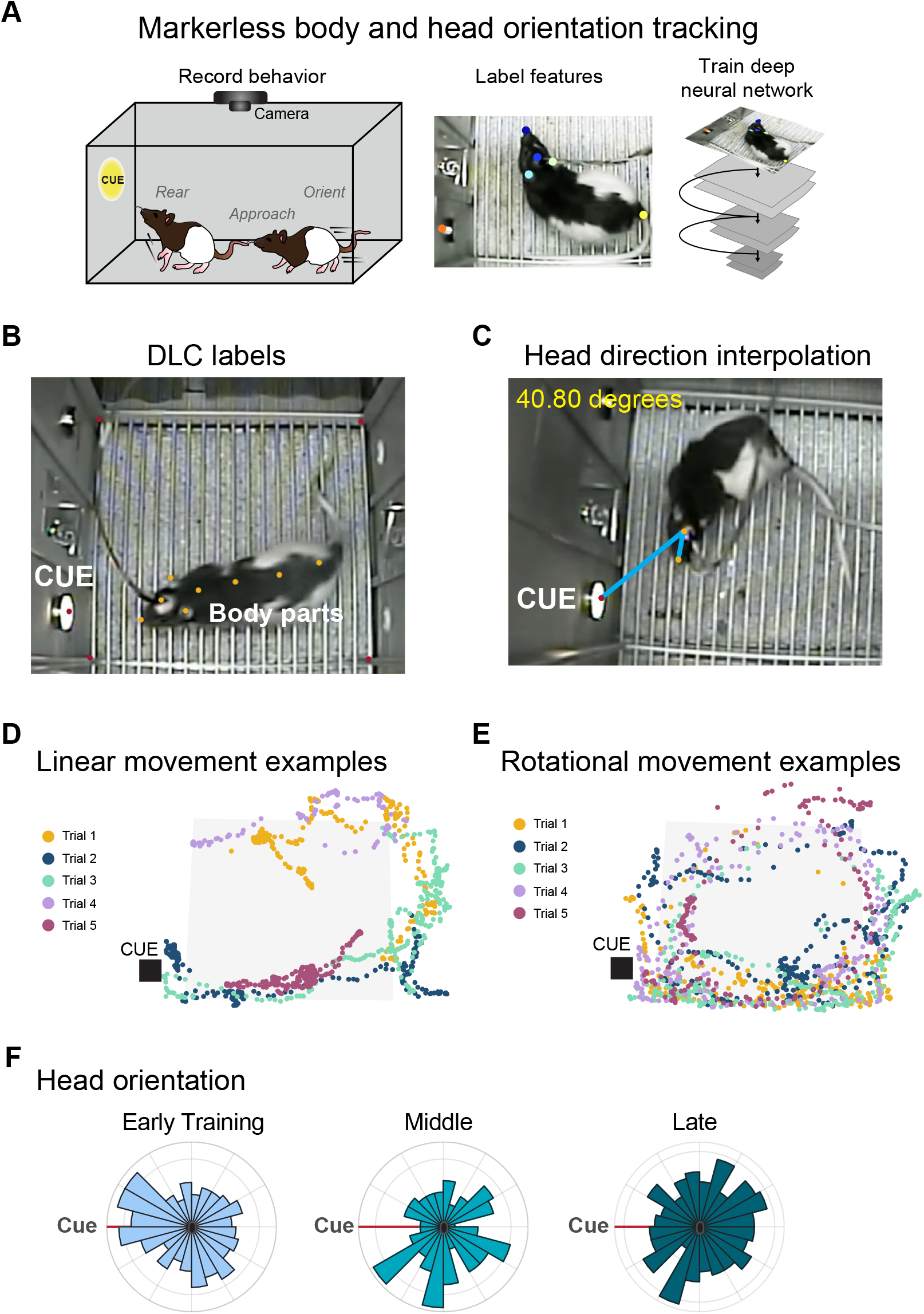
Rat pose estimation and movement analysis. A) Behavior data was extracted from video recordings using the DeepLabCut pipeline. B) Video frames containing experimenter labeled rat body parts (ears, nose, tail base, etc) and static features of the environment (cue, chamber corners), were used to train a deep learning network to estimate frame by frame x,y coordinates for the entire video data set. This data was used to calculate movement and position information across optogenetic Pavlovian conditioning. C) Head direction was interpolated by calculating the angle between the vector projecting from the cranial implant to the nose with the vector projecting from the cranial implant to the cue. D) Example trials showing linear cue-evoked movement. Each dot represents the position of the rat’s nose on an individual video frame, on 5 different trials early in training. E) Example trials showing rotational movement during 5 trials late in training. F) Polar plots showing distribution of head direction angles for an example rat across training.

**Supplemental Figure 2.**
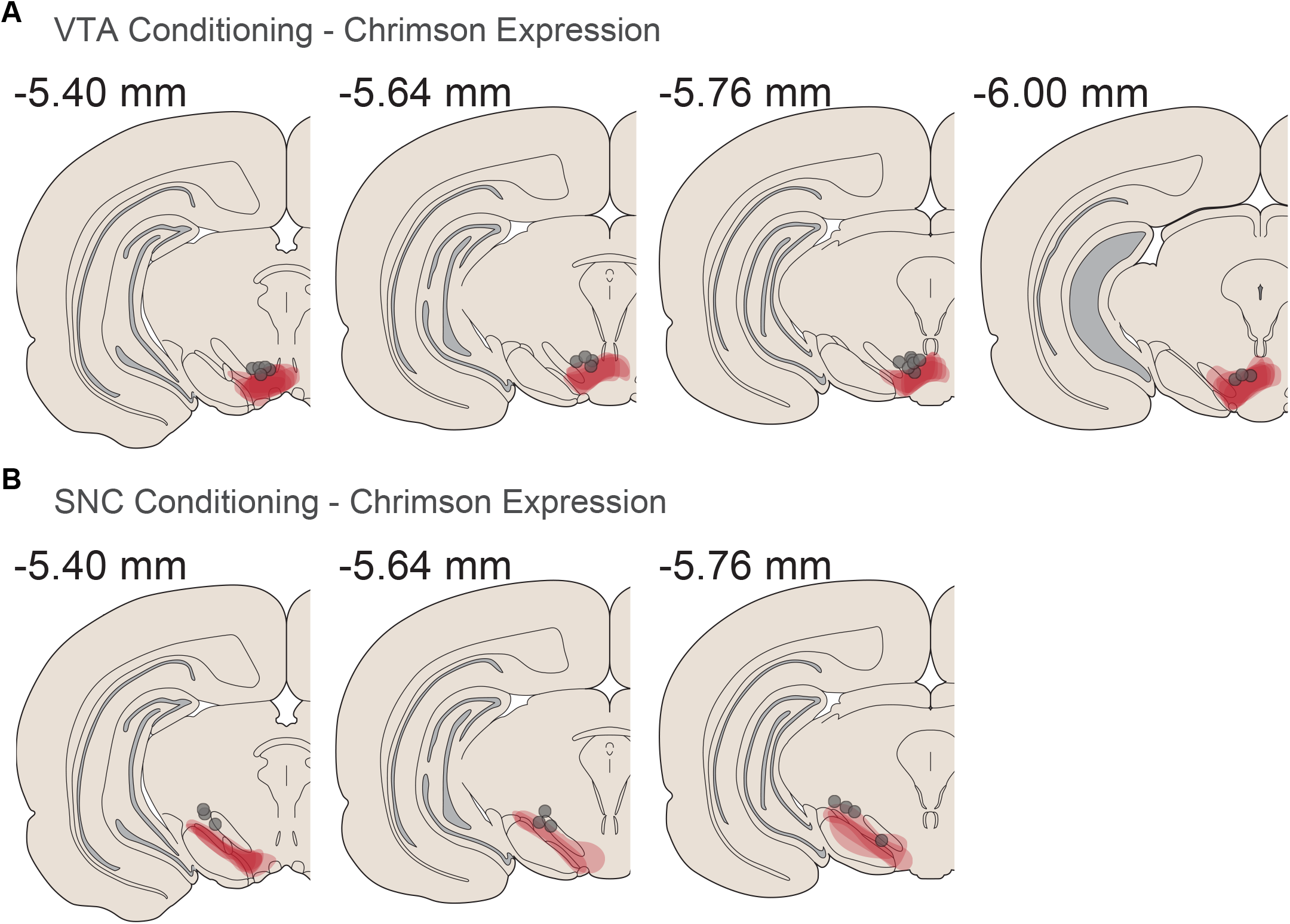
Midbrain virus expression across behavioral cohorts. A) Summary of virus expression located underneath fiber optic placements in rats in the VTA dopamine conditioning experiments. B) Summary of virus expression located underneath fiber optic placements in rats in the SNC dopamine conditioning experiments. Coordinates represent distance posterior to Bregma in the coronal plane.

**Supplemental Figure 3.**
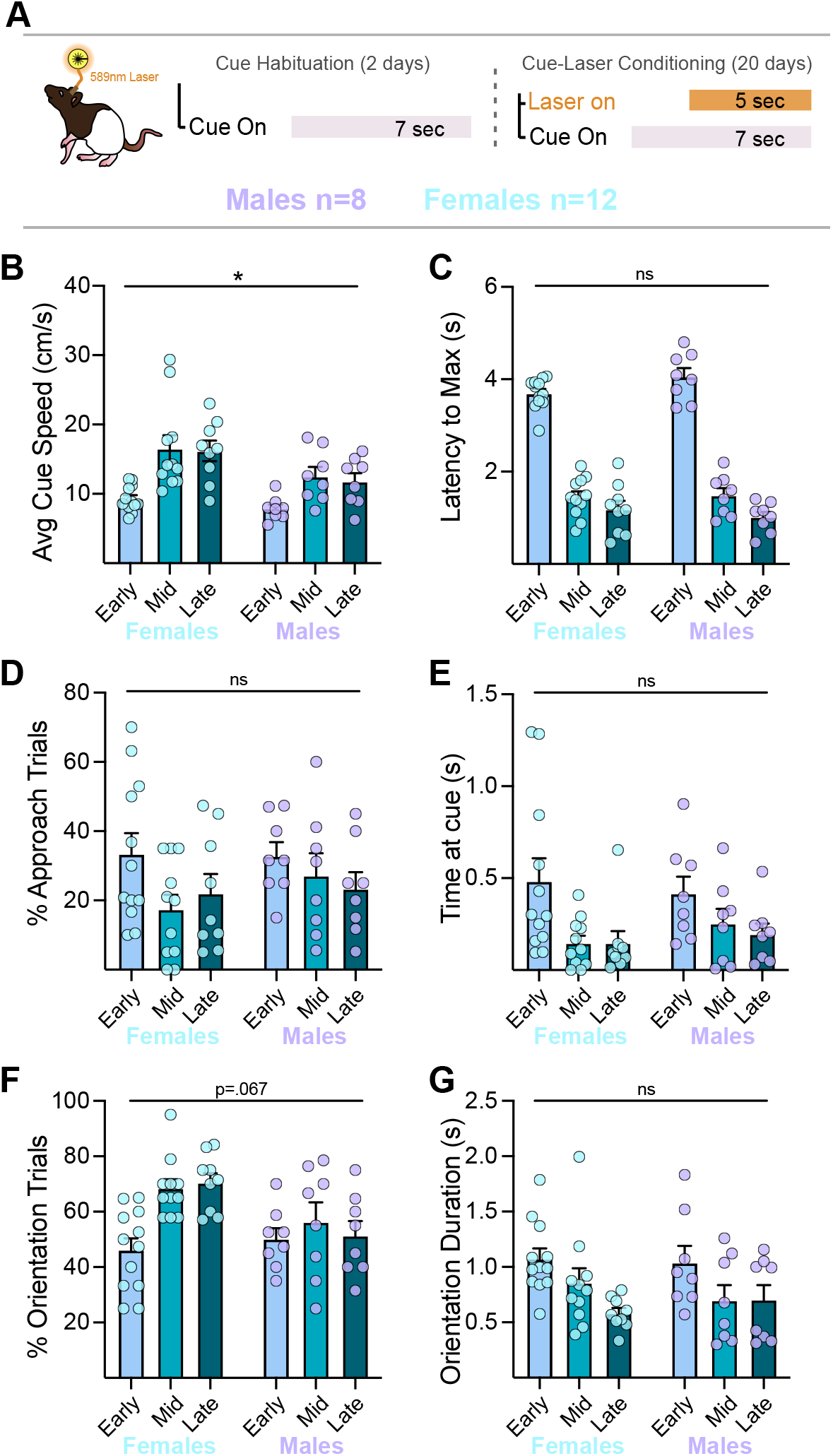
Behavioral comparisons by sex. A) Behavior during optogenetic Pavlovian conditioning was analyzed as a function of rat sex. B) Females (n=12) had higher average movement speed during cue presentations, compared to males (n=8), at all training phases. C-G) We found no other sex differences in behavioral measures, including the time to maximum movement speed, cue approach, or cue orientation. *p<.05.

**Supplemental Figure 4.**
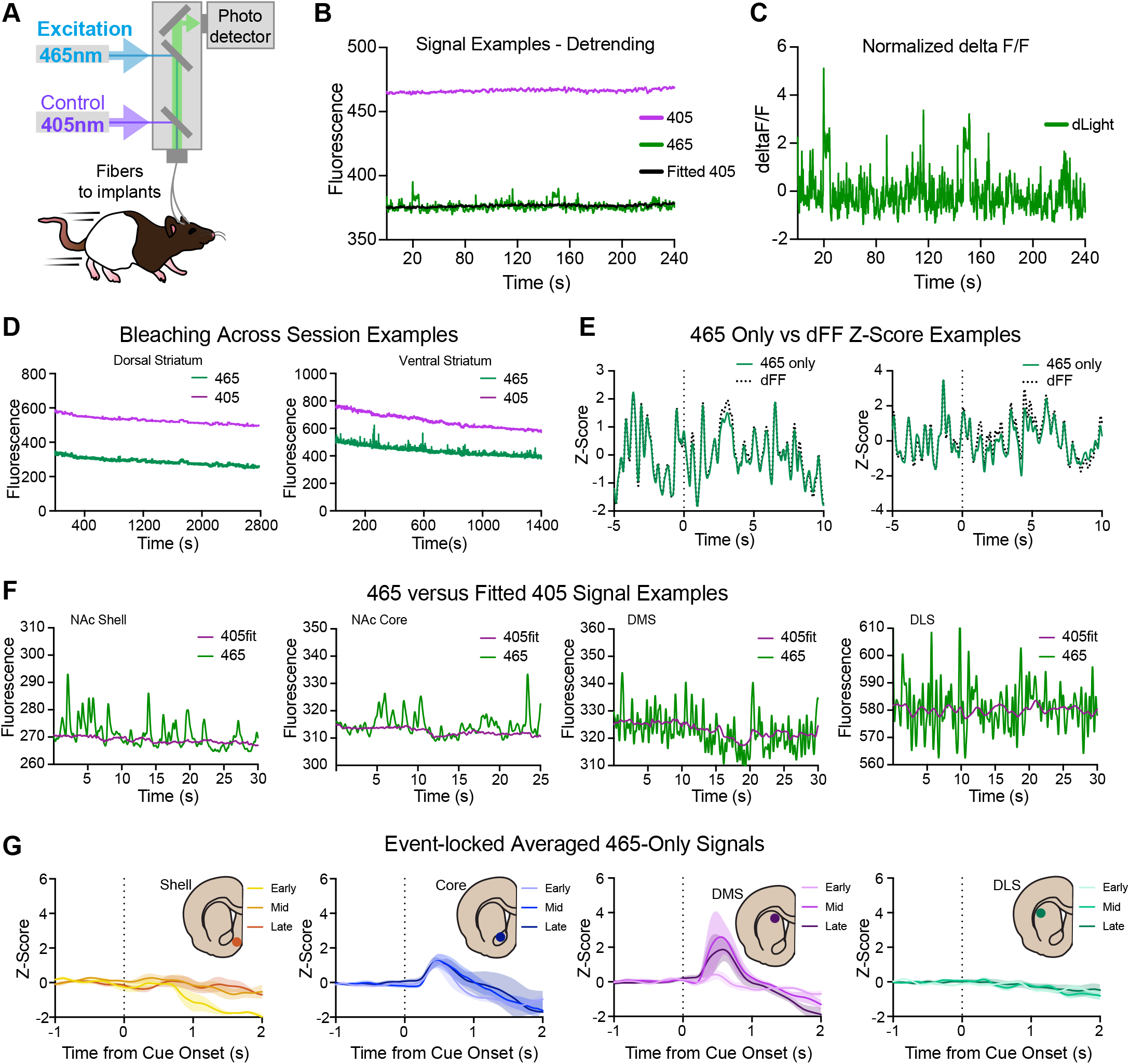
Fiber photometry analysis and signal comparisons. A) Fiber photometry set up for in vivo measurement of dopamine-related GFP fluorescence from the dLight 1.3b sensor across striatal subregions during behavioral training. All signals are first lowpass filtered (2Hz) and downsampled to 40Hz. B) Example traces of demodulated 465-nm and 405-nm signals, and the 405-nm signal after applying a least squares fit (Fitted 405). C) Normalized 465-nm trace (deltaF/F) = (465 signal - fitted 405 signal)/(fitted 405 signal). D) Example long recording traces of 405 and 465 signals show consistent bleaching patterns across signal type and region. E) Example comparisons of Z-scored 465 signal alone versus the Z-scored dFF signal calculated using the 465 and fitted 405 signals for a DMS (left) and shell (right) recording. H) Examples of 465 and fitted 405 nm signals from each region, showing high signal to noise in 465 versus 405 channels. G) Full dataset group averages of the Z-scored cue-evoked 465 signals alone show qualitatively similar patterns across training phases and regions compared to the dFF signals calculated using the 465 and fitted 405 signals (Figure 2, VTA conditioning data).

**Supplemental Figure 5.**
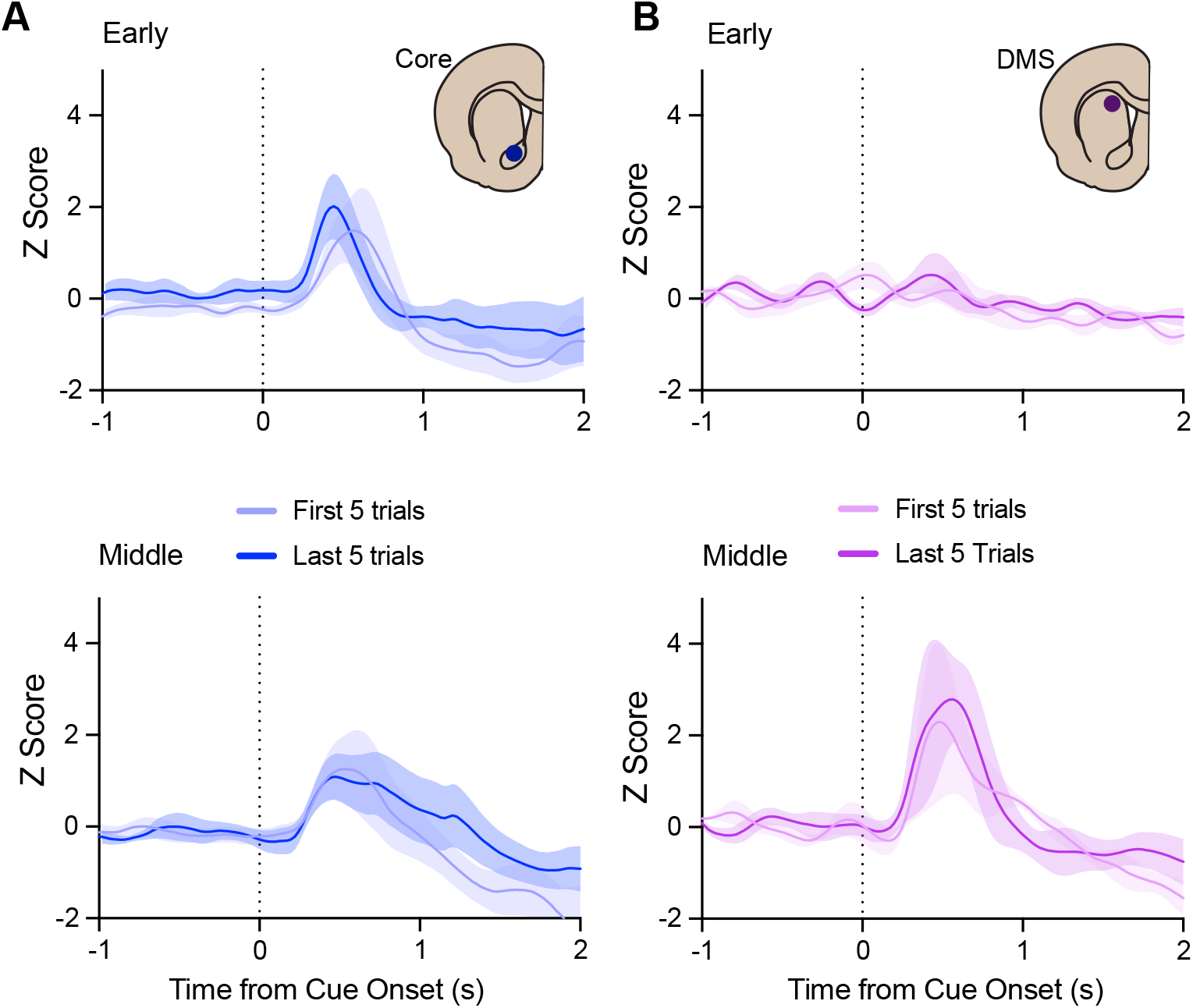
Within-session comparison of cue responses in the NAc core and DMS. dLight cue responses in the first 5 versus last five trials of the early and middle training phases are plotted for the A) NAc core and B) DMS.

**Supplemental Figure 6.**
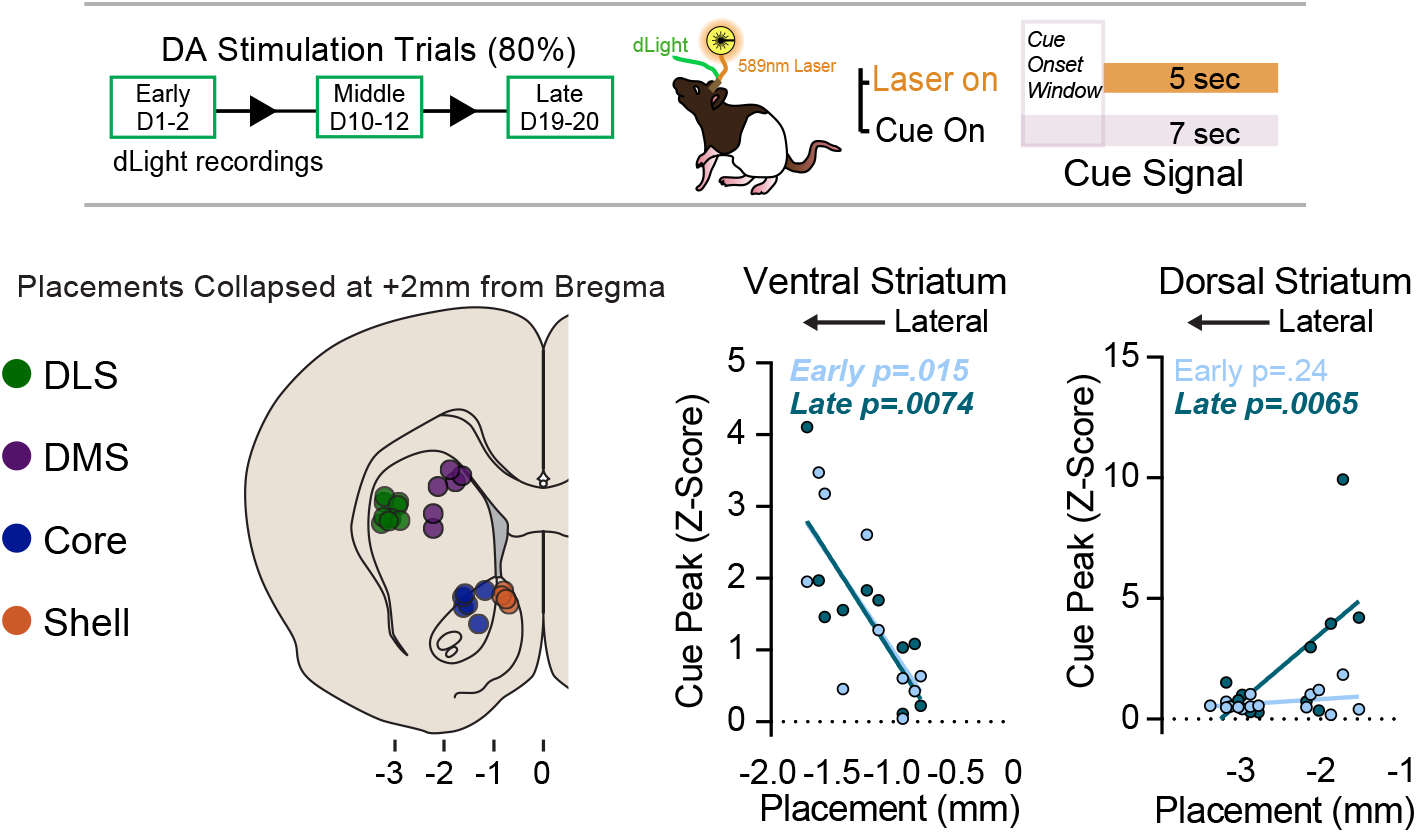
Opposing gradients of cue-evoked dopamine signals in the ventral and dorsal striatum. Anatomical placements for all recording sites were collapsed onto a coronal atlas plate at +2.0mm from Bregma. The peak cue-evoked Z-scored dLight signal at each recording site was plotted against anatomical placement on the medial to lateral axis from Bregma. Opposing placement-signal relationships were found in the ventral (NAc core and shell) and dorsal striatum (DLS and DMS) after conditioning.

**Supplemental Figure 7.**
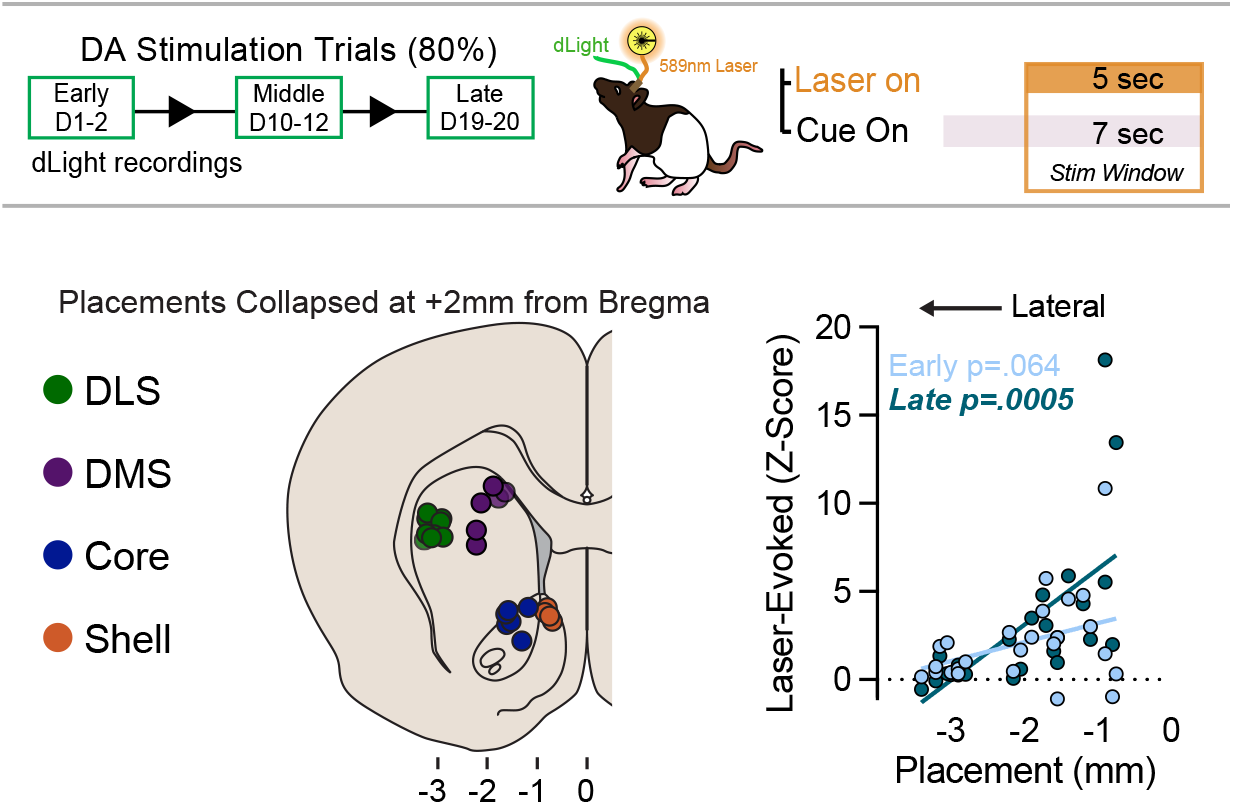
A ventrolateral to dorsomedial striatal gradient of VTA dopamine neuron evoked dopamine signaling. Anatomical placements for all recording sites were collapsed onto a coronal atlas plate at +2.0mm from Bregma. The average Z-scored dLight signal during the laser stimulation periods at each recording site was plotted against anatomical placement on the medial to lateral axis from Bregma.

**Supplemental Figure 8.**
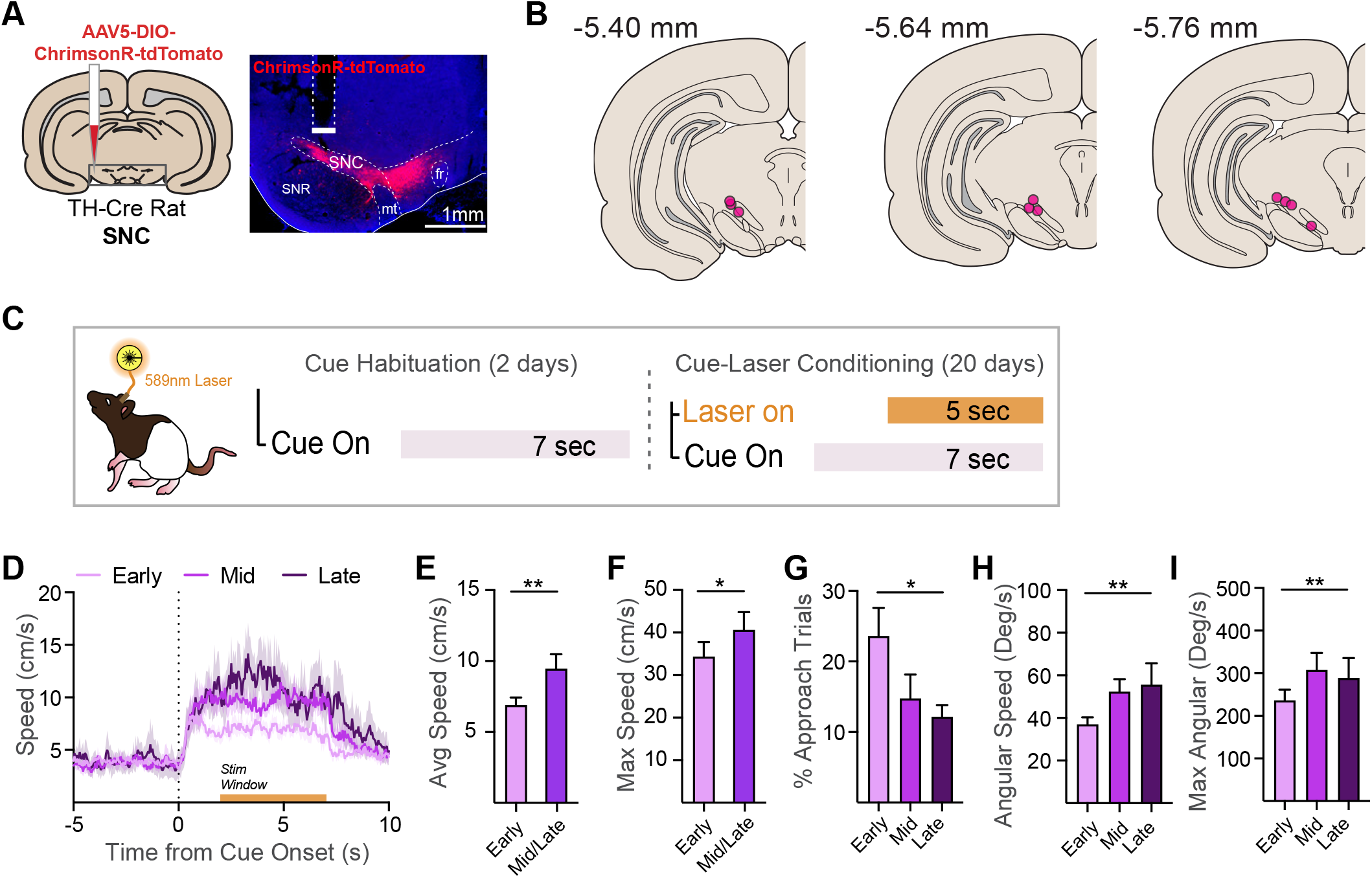
SNC dopamine neurons drive cue learning and vigorous behavior that evolves over time. A) ChrimsonR-tdTomato was expressed in dopamine neurons in TH-cre rats (N=10) and optic fibers were inserted over viral expression in the SNC. B) Location of fiber tips in the SNC. C) Schematic of optogenetic Pavlovian conditioning procedure. Following habituation to a novel, neutral cue (light + tone), rats underwent training where the cue was paired with laser (589 nm) delivery for 20 sessions. D) Across this training, conditioned responses (movement) emerged with high probability in the first 2-sec of the cue period, before laser onset. Conditioned movement was tightly locked to cue onset, increasing in speed across the early (days 1-2), middle (days 10-12), and late (days 19-20) phases of training. Both average E) and maximum F) speed achieved increased between early and middle/ late training phases. G) We also assessed the path of movement patterns, finding that rats emitted conditioned behavioral responses directed at the cue. Further classification using semi-automated pose estimation data showed a decrease in the percentage of trials containing an approach response across training. H-I) As training progressed, cue-evoked movement evolved into a more vigorous, rotational pattern, as rats ran in circles around the chamber. Rotation did not occur early in training but increased robustly in number and probability across training. Correspondingly, the H) average and I) maximum angular speed during cue presentations increased robustly across training. **p<.01, *p<.05.

**Supplemental Figure 9.**
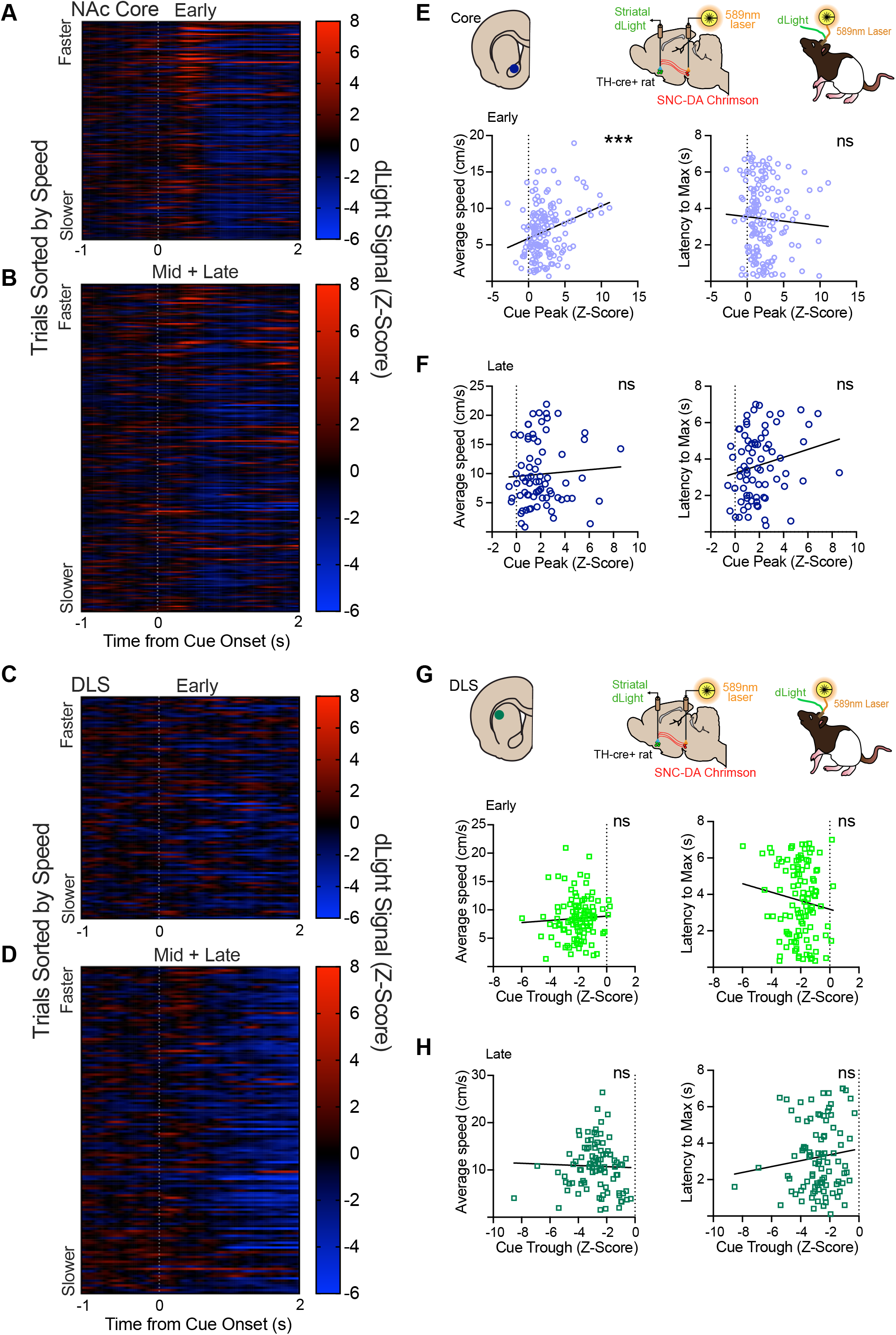
SNC cue-evoked dopamine signals are not reliably correlated with conditioned movement invigoration. A) Heatmap of Z-scored NAc core dLight signals for the early versus B) mid/late training phases centered around onset of cues predicting SNC dopamine neuron activation. Each row represents a trial, which are sorted as a function of the average speed reached during each trial, with faster movement trials on top and slower movement trials on the bottom. Corresponding heatmaps for C) early and D) mid/late DLS cue signals. E-H) Pearson correlations between the peak cue-evoked signal in the NAc core and the average speed, and latency to maximum speed, trial by trial, for early and late training phases. Cue-evoked dLight in the core was positively correlated with average speed E) early in training, but this relationship went away F) as conditioning progressed. G,H) Cue trough versus speed variable correlations for the DLS, where no significant relationships were found at any training phase. ***p<.001.

**Supplemental Figure 10.**
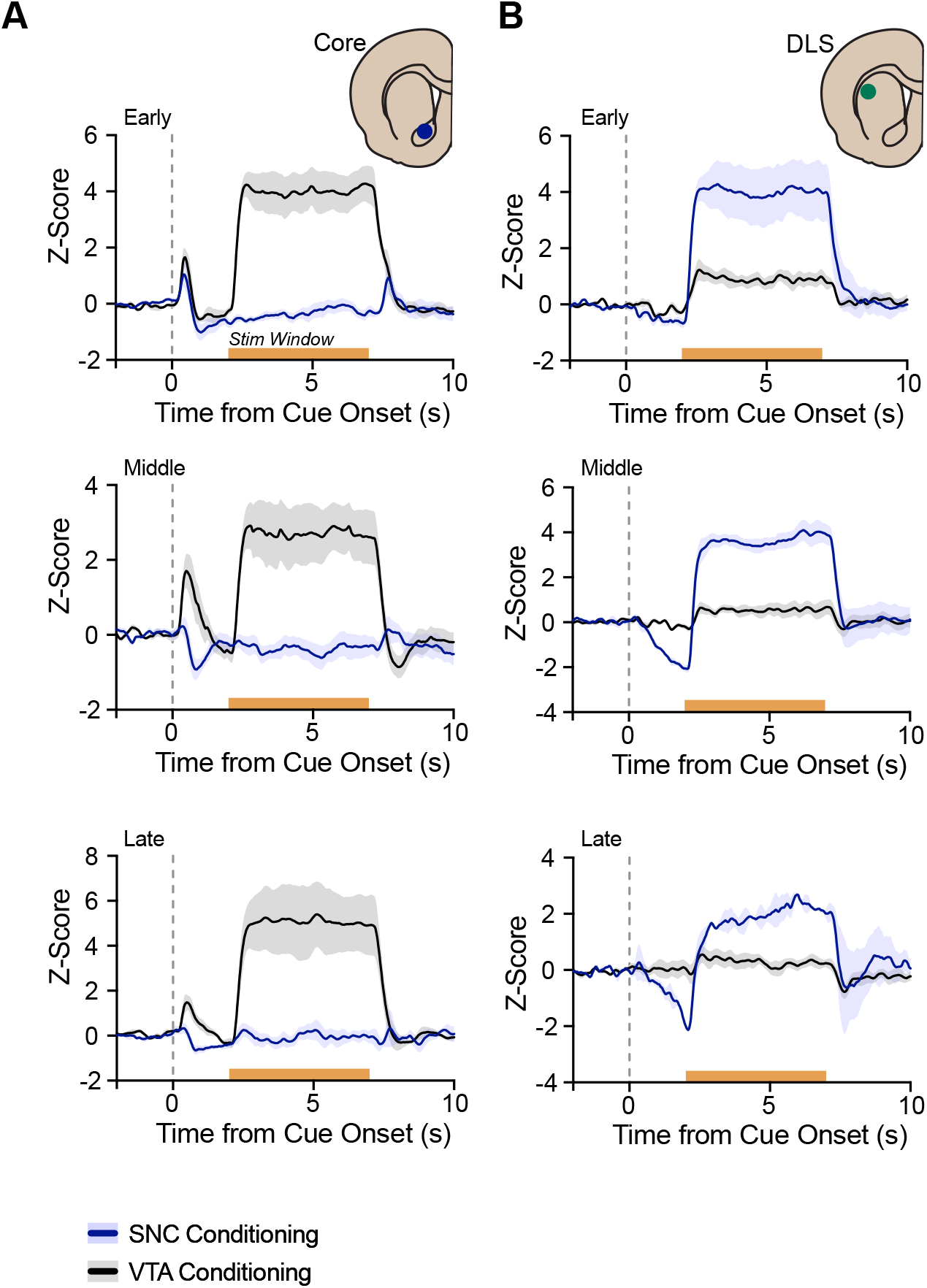
Divergent dopamine in the NAc core and DLS emerge during learning driven by VTA versus SNC dopamine neuron stimulation. A) In the NAc core, dopamine signals emerged in response to a cue predicting VTA dopamine neuron stimulation, and VTA stimulation, across optogenetic conditioning. In contrast, neither cue nor laser evoked dopamine signals were evident in the core during SNC dopamine neuron optogenetic conditioning. B) In the DLS, dopamine signals did not emerge in response to a cue predicting VTA dopamine neuron stimulation, and minimal laser-evoked signals emerged for VTA stimulation. In contrast, a decrease in dopamine in the DLS emerged to cues predicting SNC dopamine neuron stimulation, and a robust increase in dopamine in the DLS was evoked by SNC stimulation.

## Notes

### Competing Interest Statement

The authors have declared no competing interest.

### Summary of Updates

Major revisions: New data (Figure 3 and Figure 6, Suppl Figure 8,9) and several new analyses (Suppl Figures 4, 5, 10) Minor: Updated text/discussion, references incorporating new data

